# Supramolecular Enzyme-Peptide Gels for Localized Therapeutic Biocatalysis

**DOI:** 10.1101/2024.09.06.611628

**Authors:** Madeline J Fuchs, Lucas Melgar, Bethsymarie Soto Morales, Dillon T Seroski, Jennifer A Simonovich, Isabella Pinto, Chen Lu, Gianna Scibilio, Renjie Liu, Abigail Ziegler, Juanpablo Olguin, Arun Wanchoo, Ayumi Shigemoto, Dorina Avram, Benjamin G Keselowsky, Gregory A Hudalla

## Abstract

Enzyme therapeutics are often compromised by poor accumulation within target tissues, necessitating repeated systemic dosing that can exacerbate side-effects like immunogenicity. We report an injectable supramolecular enzyme-peptide gel platform for prolonged local enzyme retention in vivo. The gel is based on CATCH(+/-) (<u>C</u>o-<u>A</u>ssembling <u>T</u>ags based on <u>CH</u>arge-complementarity), cationic and anionic peptide pairs that form β-sheet fibrillar gels upon mixing. Five disparate enzymes recombinantly fused to CATCH(-) peptide integrate into CATCH(+/-) fibrils during assembly, resulting in five single-enzyme and three dual-enzyme catalytically-active enzyme-peptide gels. CATCH gels enable titratable catalytic activity and prolonged localized retention. Gels of uricase, an FDA-approved but systemically-ineffective enzyme therapy for gout, rapidly dissolve gout crystals, with low risk of anaphylaxis after repeat dosing. Dual-enzyme gels of uricase and anti-inflammatory indoleamine 2,3-dioxygenase outperformed either single-enzyme formulation by concomitantly dissolving the crystals and suppressing pro-inflammatory neutrophil functions. Modular and tunable enzyme content, coupled with prolonged retention, establish CATCH enzyme-peptide gels as a generalizable vehicle for effective local therapeutic biocatalysis.

## 1. Introduction

Harnessing enzymes as therapeutics has faced significant challenges since the mid-twentieth century. Currently there are a modest ∼40 FDA-approved enzyme-based therapeutics, which pales in comparison to the ∼120 approved antibody-based therapeutics.^1,2^ Ideally, an effective enzyme persists long enough in the appropriate tissue to exert its catalytic effector functions. For indications requiring rapid enzyme function in blood (e.g., Adzynma^®^ for blood clot prevention^3^, Idefirix^®^ for antibody eradication for transplant tolerance^4^) systemic infusion is an effective strategy. For indications requiring prolonged enzyme pharmacokinetics, chemical modifications (e.g., PEGylation, Fc-fusion) can address rapid clearance from circulation. As these modifications provide only limited accumulation in solid tissues^2,5,6^, targeting strategies, such as glycosylation in Cerezyme^®7^ and ADEPT ^8^ (Antibody-Directed Enzyme Prodrug Therapy), can be effective when unique receptor-ligand complexes exist. In the absence of such targets, localized delivery would benefit from an easily administered vehicle that is readily amenable to carrying one or more different enzymes at independently controllable doses to more effectively treat complex diseases with multiple drivers.

Biomaterials created from self-assembling peptide subunits fused to folded proteins can afford modularity and precise control of combinations of functionalities^9–12^. However, few have been designed to deliver therapeutic enzymes. We previously reported pairs of cationic and anionic β-sheet fibrillizing peptides known as CATCH(+/-) (Co-Assembly Tags based on CHarge complementarity) to create supramolecular protein-peptide gels^13^. CATCH gels are injectable, induce weak rapidly resolving inflammation, and are not immunogenic in pre-clinical models^14^. A benefit of the CATCH system is that separately, CATCH(+) and CATCH(-) peptides do not self-assemble due to electrostatic repulsion, but once combined, they co-assemble into β-sheet fibrils. As a result, either CATCH peptide can be recombinantly fused onto a larger folded protein domain, and this CATCH-tagged fusion protein can be expressed and recovered from living hosts in the soluble phase^13^. Assembly with integration of the functional domain is then triggered by simple mixing of CATCH(+), CATCH(-), and CATCH fusion protein, as shown with CATCH-Green Fluorescent Protein (GFP)^13^.

Here we introduce supramolecular enzyme-peptide gels built using the CATCH platform for localized therapeutic biocatalysis. Gels of different model enzymes demonstrated tunable activity in vitro and in vivo with both single-enzyme and dual-enzyme formulations. Gels of uricase that catabolize uric acid to allantoin showed effectiveness for locally treating gout flare by resolving uric acid crystals while not raising neutralizing nor pro-anaphylactic anti-drug antibodies. Gels of candidate anti-inflammatory enzymes, adenosine synthase A (AdsA) and indoleamine 2,3-dioxygenase (IDO), showed effectiveness in neutrophil-dependent inflammation models as single-enzyme formulations. A dual-enzyme formulation of uricase and IDO improved treatment of gout flare by enabling concurrent crystal dissolution and suppression of neutrophil activity. These data establish CATCH gels as local carriers that endow new capabilities to enzyme therapeutics beyond those that are possible with existing systemic and targeted strategies.

## 2. Results

### 2.1. CATCH enzyme-peptide gels provide tunability and prolonged local biocatalysis in vivo

CATCH-cutinase and CATCH-NanoLuc^®^ (CATCH-NL) luciferase fusions demonstrated the modular and tunable features of CATCH-enzyme peptide gels (**Figure 1**). Both fusions were expressed and recovered from *Escherichia coli* (*E. coli*) (Supplementary Fig. 1a-c). Mixtures of CATCH(+), CATCH(-), and CATCH-cutinase formed gels that converted colorless *p*-nitrophenyl butyrate (pNPB) to yellow *p*-nitrophenol (Figure 1a-b). CATCH-cutinase gel activity followed Michaelis-Menten kinetics (Figure 1c), could be titrated over two orders of magnitude (Figure 1d), and had no effect on gel mechanics (Supplementary Fig. 1d,e).

**Figure 1:**
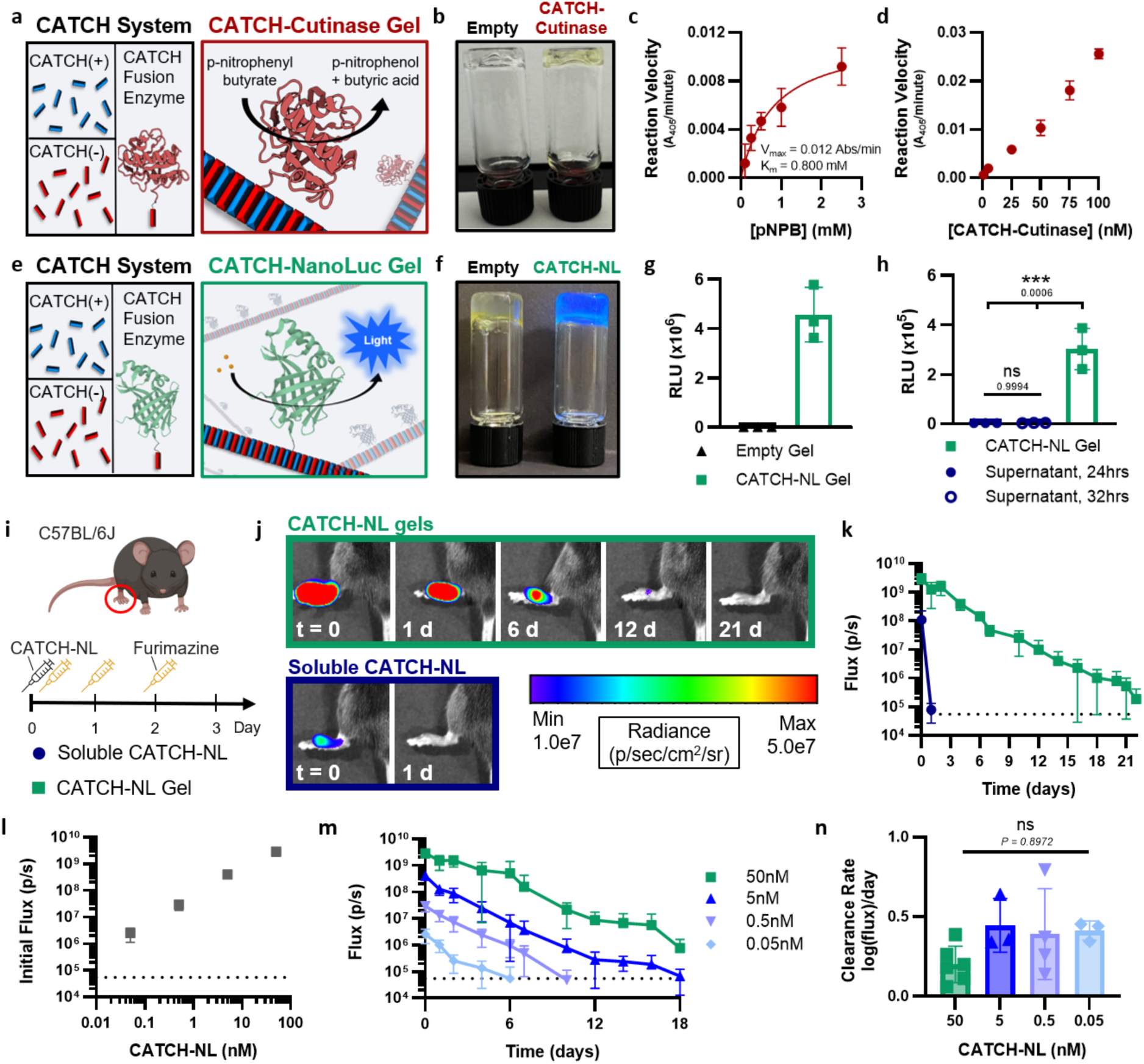
CATCH gels provide titratable and prolonged local biocatalysis. (a) Schematic of CATCH peptides and CATCH-cutinase bearing β-sheet fibrils. (b) Digital images of empty and CATCH-cutinase gels incubated with p-nitrophenyl butyrate (pNPB). (c) Reaction velocity of CATCH-cutinase gels as a function of substrate concentration. Solid line in denotes Michaelis-Menten fit of enzyme activity, N=3, R^2^=0.89. (d) Reaction velocity of CATCH-cutinase gels as a function of immobilized enzyme concentrations (N=3). (e) Schematic of β-sheet fibrils bearing CATCH-NL, catabolizing furimazine substrate (orange circles). (f) Digital images and (g) luminescence measurements of empty and CATCH-NL gels incubated with furimazine. (h) Luminescence output of PBS supernatants collected 24 and 32 h after incubation with CATCH-NL gels compared to luminescence output from the washed gels, N=3, ANOVA with Tukey’s post hoc analysis. (i) Schedule to evaluate dosing and retention of CATCH-NL subcutaneously injected into the top of the hind paw (black syringe) at day 0 and subsequent administration of furimazine over time (orange syringe), with IVIS measurements following each injection. (j) Representative IVIS images and (k) quantitative analysis of longitudinal NL activity following furimazine injection in mice that received soluble CATCH-NL (dark blue circles) or CATCH-NL gels (green squares) (dashed line represents average background), N=4. (l) Initial and (m) longitudinal flux following furimazine injections in mice that received gels formulated with varying concentrations of CATCH-NL (dashed line represents average background), N=3-6. (n) Decay rate of CATCH-NL signal over time as determined by semi-log regression (Y=10^mx+b^), N=3-6, extra-sum-of-squares F test comparing slopes across CATCH-NL concentrations, F=0.1985, average R^2^=0.943. *P ≤ 0.05, **P ≤ 0.01 ***P ≤ 0.001, ****P ≤ 0.0001. All data are presented as mean ± s.d.

The mixture of CATCH(+), CATCH(-), and CATCH-NL formed gels that bioluminesced, while gels lacking enzyme did not (Figure 1e-h). Shortly after subcutaneous injection, greater NL activity was observed for gels than soluble CATCH-NL, with CATCH-NL gel activity persisting for up to 21 days versus less than 1 day for soluble CATCH-NL (Figure 1i-k). Injection site CATCH-NL activity scaled with the molar ratio of CATCH-NL admixed with CATCH(+) and CATCH(-) during gel fabrication (Figure 1l-m). CATCH-NL activity decay rate was independent of dosing (Figure 1n), suggesting that CATCH-NL was cleared as the gel was cleared.

### 2.2. CATCH-uricase gels for localized MSU crystal dissolution

Gout is the most common form of inflammatory arthritis, affecting approximately 9.2 million adults in the U.S.^15^, where up to 35% have tophaceous disease^16^. Pegloticase, a systemic uricase variant approved by the FDA in 2010, is only used sparingly to treat gout because it shows wide variability in crystal clearance, often requiring multiple infusions over months to be effective^17,18^. A single dose of CATCH-uricase (CATCH-U) gel rapidly cleared monosodium urate (MSU) crystals at a subcutaneous gout flare site leading to diminished inflammation (**Figure 2**). CATCH-U was expressed and recovered from *E. coli* (Supplementary Fig. 2). Mixtures of CATCH(+), CATCH(-), and CATCH-U formed gels (Supplementary Fig. 3a-c) that degraded monosodium urate (MSU) crystals (Figure 2a-b) and uric acid in vitro (Figure 2c), whereas empty gels did not. At a subcutaneous injection site, CATCH-U gels had a half-life of ∼6.9 days compared to < 1 day for soluble uricase (Supplementary Fig. 3d). At a MSU crystal challenge site in C57BL/6 mice, CATCH-U gels spared initial paw distention (Figure 2d-f), more rapidly resolved paw swelling (Figure 2g, Supplementary Fig. 4), decreased IL-1β secretion (Figure 2h), and better alleviated leukocyte infiltration (Supplementary Fig. 5) compared to soluble uricase or empty gels.

**Figure 2:**
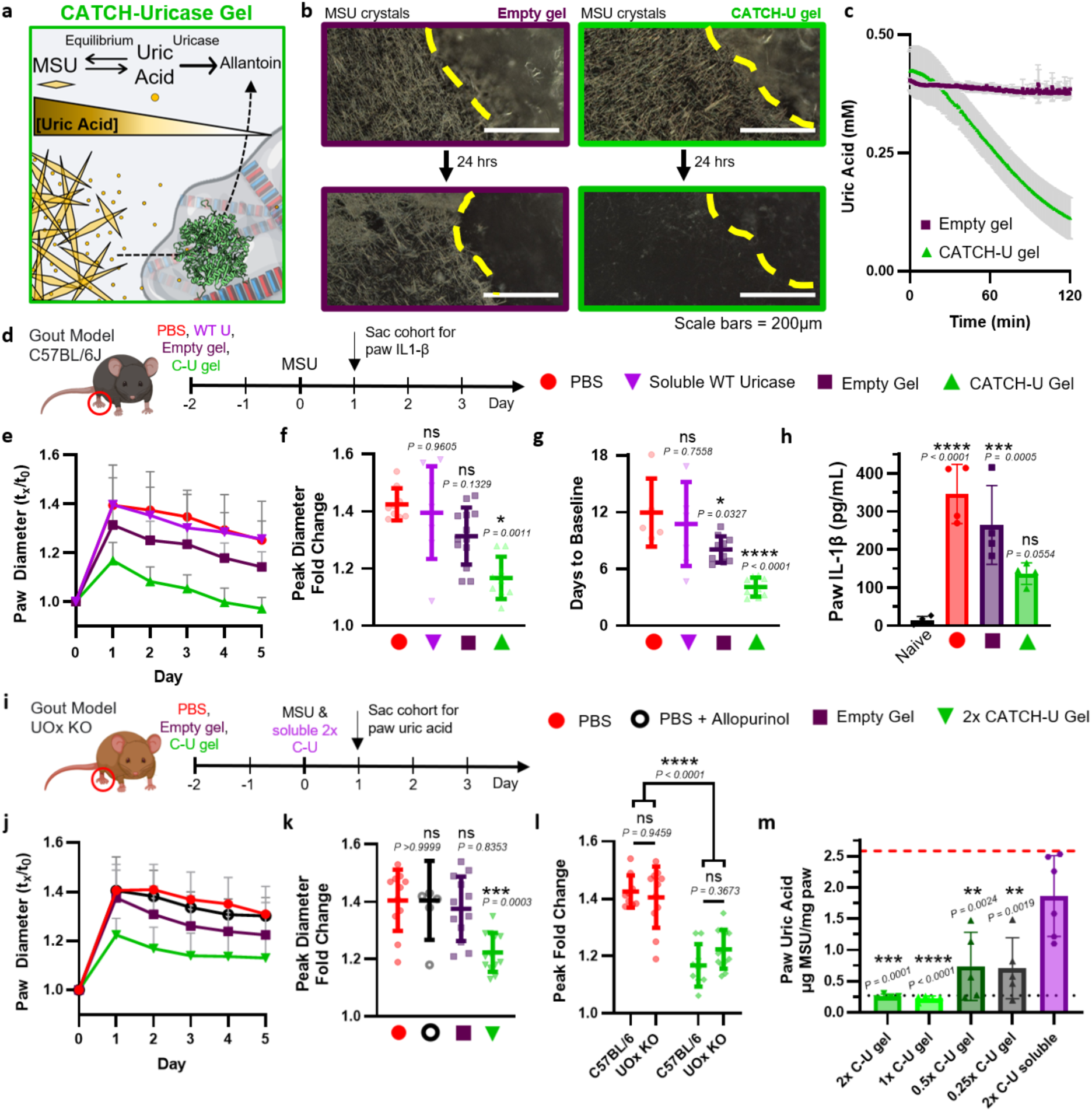
CATCH-uricase gels deplete MSU crystals. (a) Schematic of MSU crystal depletion by a CATCH-U gel, catalyzed by local enzymatic conversion of uric acid to allantoin. (b) Micrographs of empty and CATCH-U gels (gel edge indicated by dashed yellow line) incubated with MSU crystals at t = 0 and 24 h. Scale bar = 200 µm. (c) Level of uric acid present in solution containing empty (N=4) versus CATCH-U gels (N=3). (d) Schedule to evaluate inflammation following subcutaneous MSU challenge in C57BL/6J mice pretreated with PBS (red circle), soluble wild-type (WT) uricase (purple inverted triangle), empty gel (plum square), or CATCH-U gel (green triangle). (e) Fold change in paw diameter relative to day 0, (f) peak fold change, and (g) extrapolated time to baseline, average R^2^ = 0.880. N = 8-13, ANOVA with Dunnett’s multiple comparisons test to PBS group. (h) IL-1β in MSU challenged paw. N=4, ANOVA with Dunnett’s multiple comparisons test to PBS group. (i) Schedule to evaluate inflammation following subcutaneous MSU challenge in UOx KO mice pretreated with PBS (red circle), sustained on allopurinol water (black open circle), empty gel (plum square), or 2x CATCH-U gel (upside down green triangle). (j) Fold change in paw diameter relative to day 0, and (k) peak fold change. N=11-13, ANOVA with Dunnett’s multiple comparisons test to PBS group. (l) Peak fold change in paw diameter relative to day 0 for mice treated with PBS or 1x CATCH-U gel in C57BL/6 mice and PBS or 2x CATCH-U gel in UOx KO mice. N=9-13, ANOVA with Tukey’s multiple comparisons. (m) MSU crystal burden in mouse paws 24 hours after MSU injection in mice pretreated with PBS or varying concentration CATCH-U gels 48 hours prior to MSU injection, and mice treated with 2x soluble CATCH-U 1 hour after MSU injection. N=4-6 ANOVA with Dunnett’s multiple comparisons test to 2x soluble C-U group. Red dashed line (top) and black dotted line (bottom) represent average uric acid in PBS pretreated paws (N=6) and untreated paws (N=12), respectively. *P ≤ 0.05, **P ≤ 0.01 ***P ≤ 0.001, ****P ≤ 0.0001. All data are presented as mean ± s.d.

While C57BL/6 mice are a common model for gout^19–22^, they are imperfect because the mice express liver uricase that maintains low serum uric acid levels^23^. Humans lack a functional uricase gene and so are at increased risk of elevated serum uric acid levels that seed MSU crystallization^24^. Clinically, elevated serum uric acid can be reduced by the xanthine oxidase inhibitor allopurinol; however, systemic allopurinol shows limited effectiveness as a first-line therapy for gout flare due to inconsistent ability to clear crystals^25–27^. A single dose of CATCH-U gel in uricase knockout mice (UOx KO) cleared MSU crystals at a subcutaneous site more effectively than allopurinol, resulting in diminished gout flare (Figure 2i-m). In hyperuricemic UOx KO mice challenged with MSU crystals, CATCH-U gels spared initial paw distention more effectively than allopurinol treatment alone or blank gels (Figure 2j-k, Supplementary Fig. 6a,b), and to a similar extent as in C57BL/6J mice, which maintain low serum uric acid (Figure 2l, Supplementary Fig. 6c,d). Gels containing 2x to 0.25x CATCH-U cleared tissue crystal burden more effectively than soluble uricase (Figure 2m, Supplementary Fig. 6e,f), with the 0.25x CATCH-U dose of 0.16 mg/kg being comparable to the clinically used Pegloticase dose of 0.13 mg/kg^28^, assuming average human and mouse weight of 60 kg and 0.02 kg, respectively.

Notably, though, this dose of Pegloticase is typically given over a 2-hour infusion, bi-weekly for several months before significant dissolution of crystals, if any, occurs^18,28,29^. CATCH gels enable localized uricase action that provides rapid debulking of gouty tophi, directly resolving the pathological driver of acute and chronic disease^30,31^.

In the clinic, repeated dosing of Pegloticase (Krystexxa^®^) leads to the emergence of anti-drug antibodies (ADAs) that increase the incidence of infusion reaction, putting the patient at risk of anaphylaxis or other adverse immune reactions, necessitating discontinuation^18,32^. Anti-Pegloticase ADAs correlate with faster drug clearance and a loss of urate-lowering effect. Similarly, the *Aspergillus flavus* variant of uricase (marketed as Rasburicase), cloned in CATCH-U, is known to be immunogenic when delivered systemically over repeated doses.^33^ Here we used an immunogenicity safety study to assess the risk profile of CATCH-U gels (**Figure 3**). In this study, mice receive a subcutaneous injection of a 5x/2.5x dose prime-boost regimen of CATCH-U gel, CATCH-U in PBS (vehicle cohort), or CATCH-U in an experimental adjuvant (TiterMax^®^ Gold, adjuvanted cohort) on day 0 and 28 (Figure 3a).

**Figure 3:**
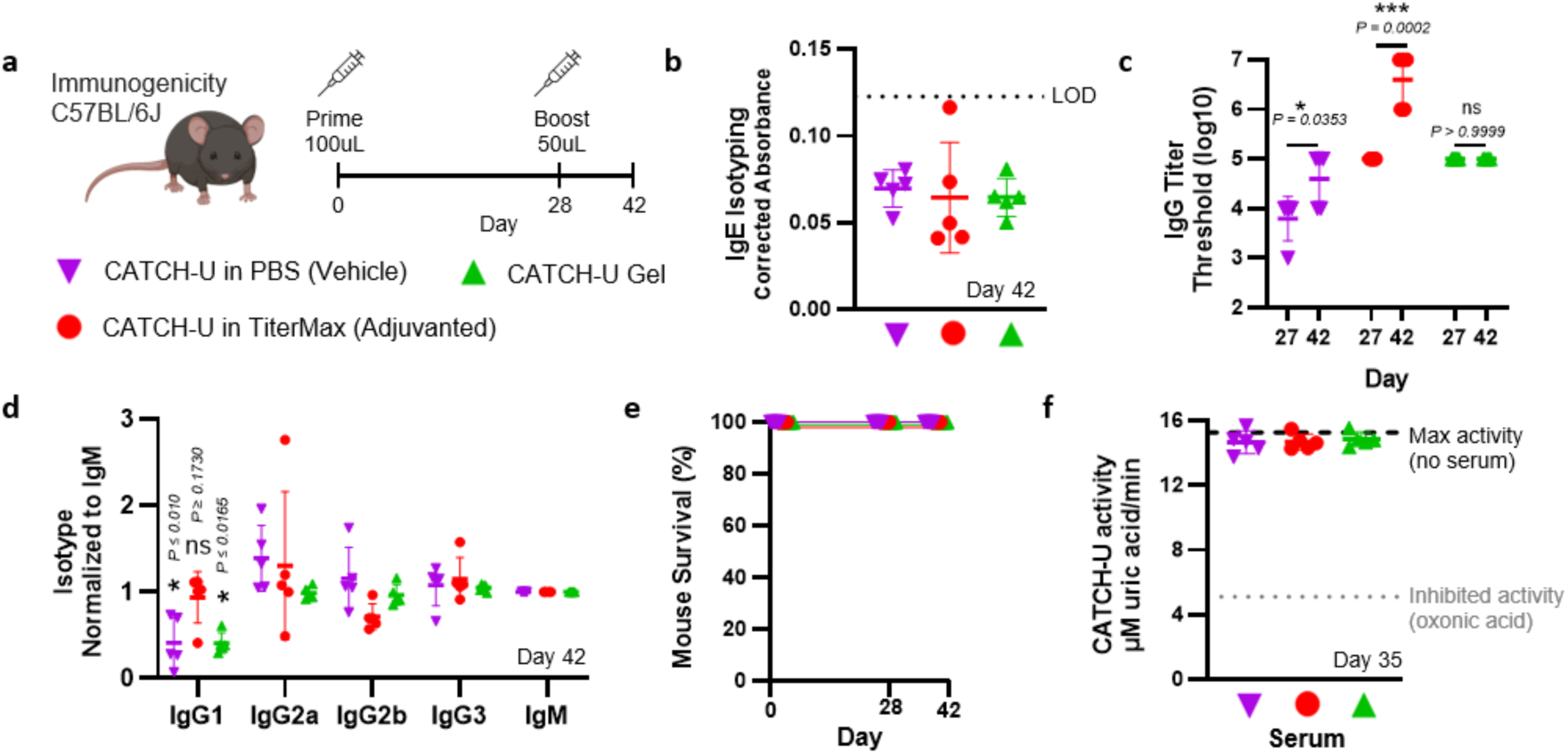
Antibodies raised to CATCH-U gels do not induce anaphylaxis nor neutralize activity. (a) Schedule to evaluate antibody formation to CATCH-U formulated in PBS (vehicle, purple inverted triangle), TiterMax (adjuvanted, red circle), or in CATCH gel (green triangle), N=5 per group. (b) IgE isotyping on day 42, with dashed line indicating limit of detection (LOD). (c) IgG titer threshold on day 27 and day 42, two-tailed t-test between timepoints within treatment group. (d) IgG isotype normalized to IgM on day 42, ANOVA within treatment group with Dunnett’s multiple comparisons test to IgG1. (e) Mouse survival over the course of experiment. (f) Enzymatic activity of CATCH-U in vitro incubated with mouse serum from day 35. *P ≤ 0.05, **P ≤ 0.01 ***P ≤ 0.001, ****P ≤ 0.0001. All data are presented as mean ± s.d.

IgE antibodies against uricase, which are associated with risk of anaphylaxis^34^, were not detected in any cohort (Figure 3b). Consistent with clinical findings^33^, C57BL/6 mice raised antibodies against CATCH-U, whether delivered in the vehicle, adjuvanted, or gel form (Figure 3c). However, while vehicle and adjuvanted formulations induced a multi-titer boost response, unexpectedly total IgG remained unchanged after the CATCH-U gel boost. Furthermore, post-boost titers for the CATCH-U gel cohort were comparable to the vehicle cohort and lower than the adjuvanted cohort (p = 0.36 and p = 0.006 respectively, ANOVA with Dunnett’s multiple comparisons test to gel cohort). This contrasts starkly with other fibrillar peptide biomaterials which robustly boosted anti-protein antibodies in other contexts.^11,35–37^ Whereas the isotype profile of the adjuvanted cohort was balanced across IgG subclasses, the CATCH-U gel cohort yielded lower IgG1 levels relative to the other subclasses, having a similar IgG isotype profile to the vehicle cohort (Figure 3d). This is again in contrast to other fibrillar peptide biomaterials which typically elicit primarily IgG1 anti-protein antibodies^11,36,37^. Low IgG1 levels are associated with lowered risk of anaphylaxis^34,38^. Indeed, all mice survived and were healthy throughout the study (Figure 3e), indicating a low anaphylaxis risk with repeat CATCH-U gel use. Finally, the ADAs raised did not neutralize uricase enzymatic activity in vitro (Figure 3f), suggesting that any anti-uricase antibodies raised are not expected to diminish drug effectiveness through drug inactivation at the gout flare site.

In this study design, animals received CATCH-U formulations containing 3.2 mg/kg (prime) plus 1.6 mg/kg (boost), assuming an average mouse weight of 0.02 kg. For context, these doses are over an order of magnitude greater than clinical uricase doses^28,39^. These data suggest intra-tissue delivery of a CATCH-U gel can shift the immunoreactivity profile away from type I hypersensitivity and anaphylaxis, instead toward a local hypersensitivity akin to the resolving Arthus reaction seen with some repeated-use vaccines^40^.

### 2.3. CATCH enzyme gels to suppress local inflammation

During acute gout flare, neutrophils swarm within minutes and are activated by MSU crystals^30,41^, driving the inflammation seen within hours and lasting for days in preclinical challenge models^22^. Clinically, oral colchicine is prescribed to suppress neutrophil function in gout, but its narrow therapeutic index creates a high risk for toxicity^42^. We adapted CATCH gels to locally suppress neutrophil activation by installing candidate anti-inflammatory enzymes adenosine synthase A (AdsA) (**Figure 4**) or indolamine 2,3-dioxygenase (IDO) (**Figure 5**). AdsA is a *Staphylococcus aureus* enzyme that acts as an immune evasion factor by catalyzing dephosphorylation of extracellular ATP, a pro-inflammatory signal, into adenosine, an anti-inflammatory signal^43,44^, and to our knowledge, has not been explored as a therapeutic before. IDO catabolizes the essential amino acid tryptophan into kynurenine and has previously been shown to be effective for local suppression and resolution of inflammation^45,46^.

**Figure 4:**
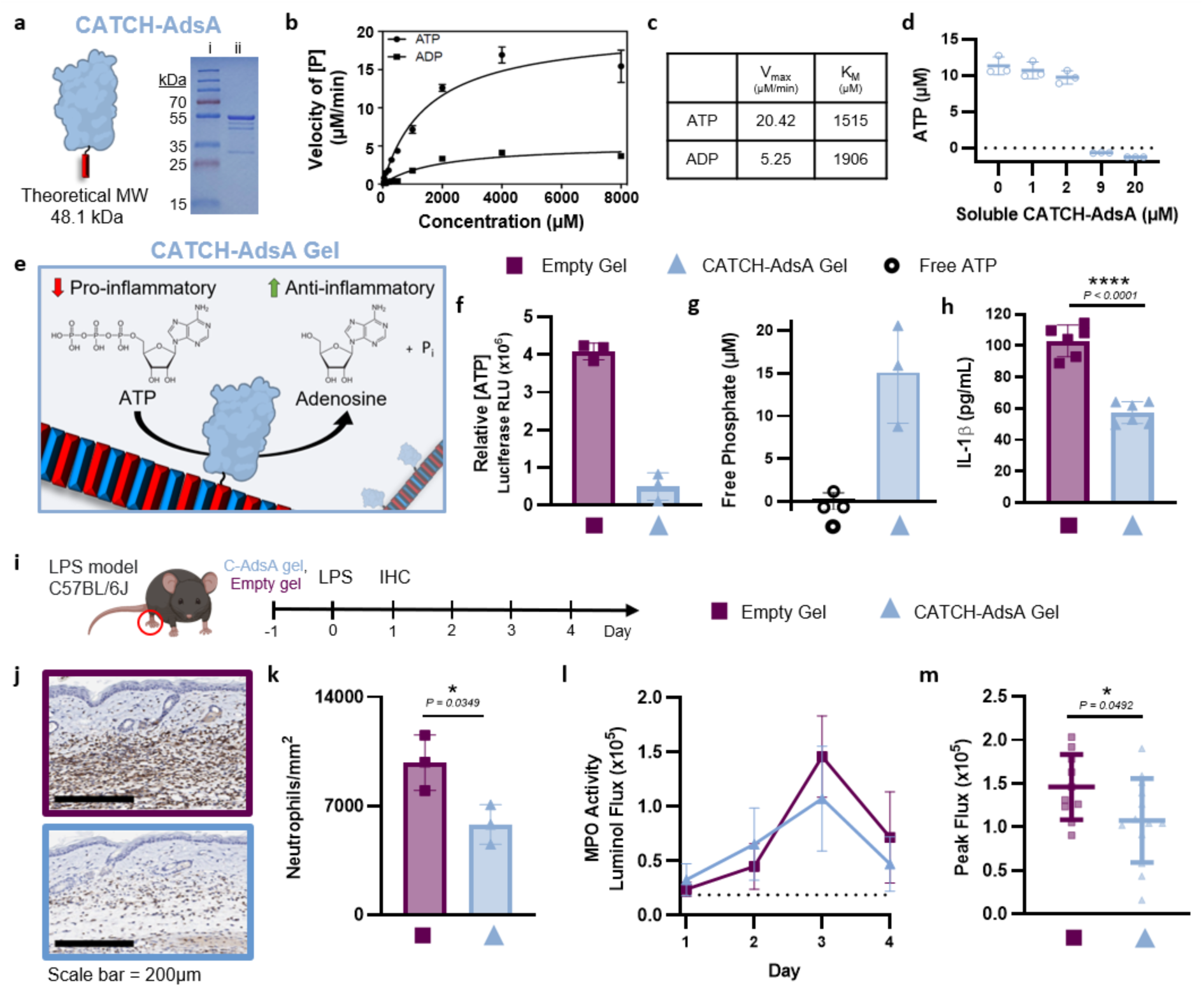
CATCH-AdsA gels deplete ATP and suppress local LPS-induced inflammation. (a) SDS PAGE gel of CATCH-AdsA purified from *E. coli*. (b) Reaction velocity of soluble CATCH-AdsA with ATP or ADP as measure by free phosphate generation ([P]) (N=3), where the solid line denotes Michaelis-Menten fit with (c) the parameters listed in the table. (d) ATP depletion by varying concentrations of soluble CATCH-AdsA, N=3. (e) Schematic of CATCH-AdsA β-sheet fibril activity. (f) Relative ATP concentration, as measured via in vitro luciferase bioluminescence, of empty and CATCH-AdsA gels incubated with ATP for one hour, N=3. (g) Free phosphates generated by ATP incubated alone or with CATCH-AdsA gels for hour, N=3. (h) IL-1β secreted from PMA-treated THP-1 cells stimulated in vitro with LPS overnight followed by ATP-supplemented media that was pre-incubated with either an empty or CATCH-AdsA gel. N=6, two-sided t-test. (i) Schedule to evaluate subcutaneous LPS challenge via daily luminol measurements or immunohistochemistry (IHC) at t = 1 d. Animals received empty gels (plum square) or 9 μM CATCH-AdsA gels (light blue triangle) on day -1. (j) Representative anti-Ly6B.2 IHC images (scale bar =200μm) and (k) neutrophil number per image area, N=3, two-sided t-test. (l) MPO activity and (m) peak flux as measured by IVIS luminol bioluminescence in mice that received empty or CATCH-AdsA gels, N = 10-13, two-sided t-test. *P ≤ 0.05, **P ≤ 0.01 ***P ≤ 0.001, ****P ≤ 0.0001. All data are presented as mean ± s.d.

**Figure 5:**
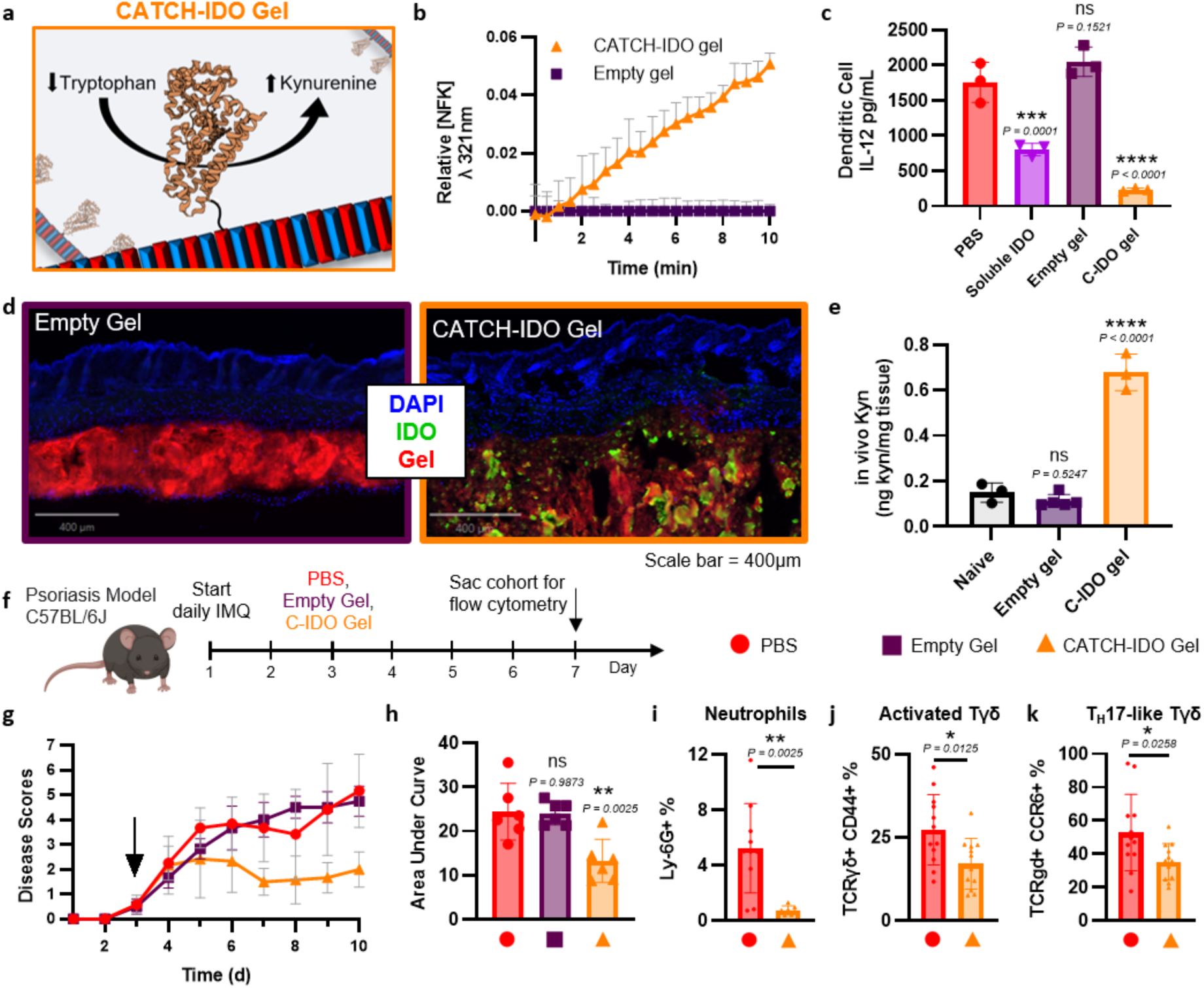
CATCH-IDO gels generate kynurenine and reduce disease severity in imiquimod-induced psoriasis. (a) Schematic of CATCH-IDO β-sheet fibrils. (b) Relative N-formyl kynurenine (NFK) concentration generated from CATCH-IDO gels normalized to empty gels, N=2. (c) IL-12 produced by LPS-stimulated mouse bone marrow derived dendritic cells pretreated overnight with PBS, soluble IDO, empty gel, or CATCH-IDO gel. N=3, ANOVA with Dunnett’s multiple comparisons to PBS group. (d) Immunofluorescent histology of mouse back skin 2 hours after injection with empty gel (left) or CATCH-IDO gel (right) (scale bar = 400μm). (e) Kynurenine (kyn) levels from mouse ear tissue 1 hour after injection with empty gel or CATCH-IDO gel compared to naive tissue. N=3-5, ANOVA with Dunnett’s multiple comparison test to naive. (f) Schedule to evaluate disease severity in imiquimod (IMQ)-induced psoriasis mice treated with PBS (red circle), empty gels (plum square), or CATCH-IDO gels (orange triangle) at first sign of symptoms (day 3), with cohorts sacrificed at day 7 for evaluation of immune cell populations. (g) Resultant clinical disease severity scores and (h) area under the curve. N=6, ANOVA with Dunnett’s multiple comparisons to PBS group. (i-k) Immune cell populations in the dermis on day 7 in mice treated with PBS or CATCH-IDO gels. N=7-12, two-tailed t test. *P ≤ 0.05, **P ≤ 0.01 ***P ≤ 0.001, ****P ≤ 0.0001. All data are presented as mean ± s.d.

CATCH-AdsA expressed and recovered from *E. coli* dephosphorylated ATP and ADP consistent with prior reports^43^ (Figure 4a-d, Supplementary Fig. 7a) and was active over a wide pH and temperature range (Supplementary Fig. 7b,c). Mixtures of CATCH(+), CATCH(-), and CATCH-AdsA formed gels that depleted ATP, thereby generating free phosphates from bulk media (Figure 4e-g), and suppressed secretion of IL-1β by PMA-differentiated THP-1 cells (Figure 4h), which is induced by extracellular ATP after LPS priming.^47^ In the neutrophil-dependent LPS inflammation model, CATCH-AdsA gels blunted Ly6-B.2+ neutrophil infiltration and degranulation more effectively than empty gels (Figure 4i-m, Supplementary Fig. 7d-f).

CATCH-IDO was expressed and recovered from *E. coli* (Supplementary Fig. 8a-d). Mixtures of CATCH(+), CATCH(-), and CATCH-IDO formed gels that generated N-formyl kynurenine in vitro (Figure 5a-b) and suppressed LPS-induced secretion of IL-12 and IL-6 from mouse bone marrow-derived dendritic cells in vitro, more effectively than soluble IDO (Figure 5c, Supplementary Fig. 8e). CATCH-IDO gels generated kynurenine in vivo at subcutaneous injection sites (Figure 5d-e, Supplementary Fig. 8f, Supplementary Fig. 9). CATCH-IDO gels suppressed the progression of IMQ-induced psoriasis (Figure 5f-h, Supplementary Fig. 10a-d), a neutrophil-driven disease^48,49^ for which localized IDO has previously been shown to be effective^46,50^. In the dermis on day 7 of imiquimod challenge, CATCH-IDO gels reduced the frequency of pathogenic Ly-6G+ neutrophils, CD44+ activated TCRγδ+ T cells, and Th17-like CCR6+ TCRγδ+ T cells (Figure 5i-k). In the draining lymph node, CATCH-IDO gel treatment decreased the frequency of CD8+ T cells and CCR6+ Th17 cells (Supplementary Fig. 10e). Finally, CATCH-IDO gels increased the frequency of immunosuppressive T regulatory cells in both the dermis and lymph nodes (Supplementary Fig. 10e,f).

### 2.5 CATCH dual-enzyme gels to suppress gouty inflammation

Gout flare is driven by local MSU crystal burden and the associated inflammation from neutrophil recruitment and activation^30,41^. Clinically, flares are treated with medications that separately modulate the uric acid burden or the inflammation, such as allopurinol and colchicine, respectively^25^. As an alternative, we evaluated dual-enzyme CATCH gels harboring CATCH-U and either CATCH-AdsA or CATCH-IDO to act as a more potent therapeutic for local gout flare than CATCH-U alone (**Figure 6**). To demonstrate feasibility of installing two enzymes, gels of CATCH-NL and CATCH-cutinase were fabricated by varying the enzyme molar ratios in solution during assembly (Figure 6a). The model dual-enzyme gels generated yellow p-nitrophenol and blue bioluminescence upon addition of pNPB and furimazine respectively (Figure 6b), with the reaction velocity of CATCH-cutinase predictably titrated into CATCH-NL gels (Figure 6c).

**Figure 6:**
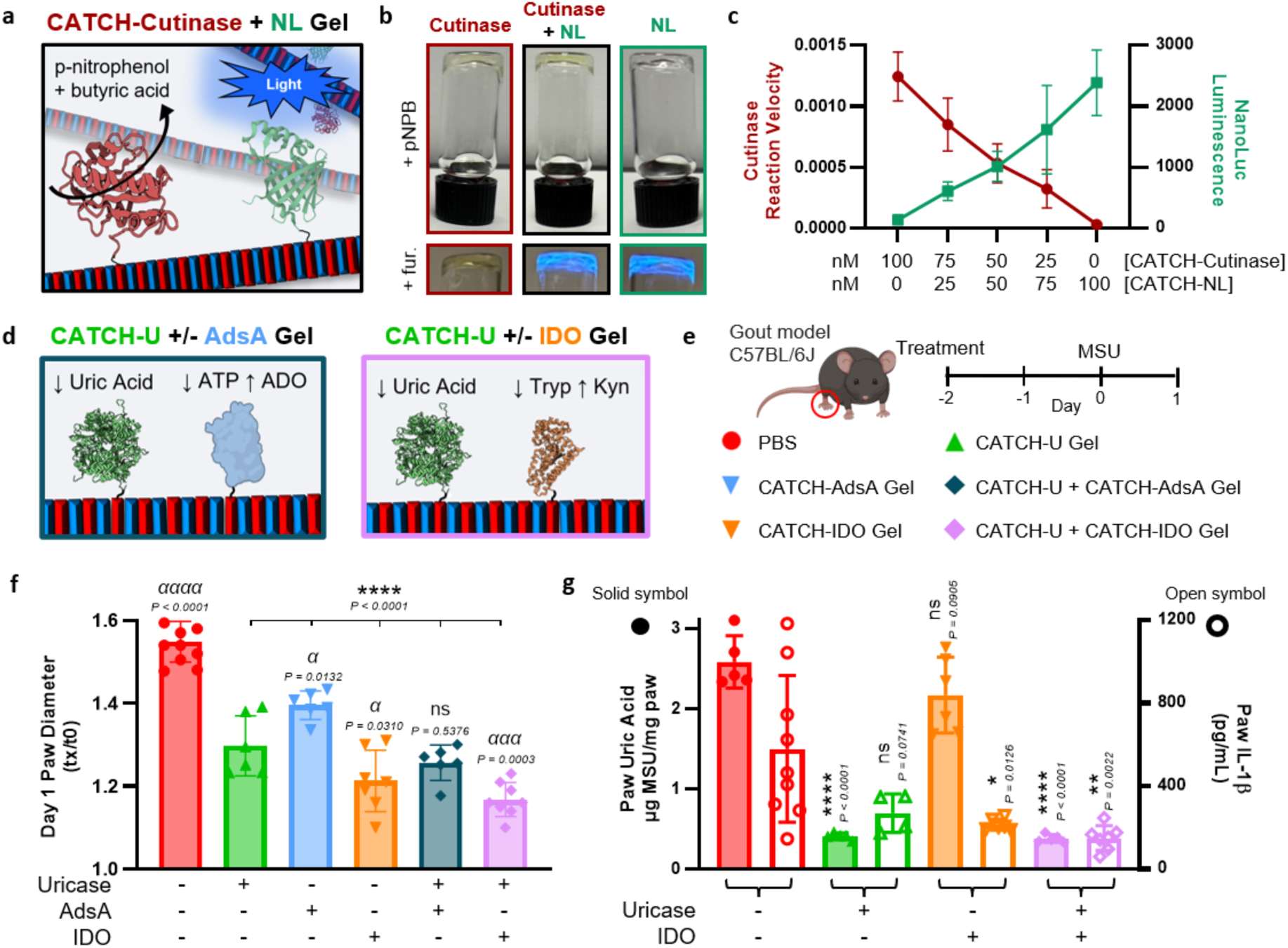
Dual-enzyme gels are titratable and more effectively reduce inflammation in gout. (a) Schematic of CATCH-cutinase and CATCH-NL dual-enzyme β-sheet fibrils. (b) Digital images of a CATCH-cutinase gel (left), CATCH-cutinase + CATCH-NL dual-enzyme gel (middle), and CATCH-NL gel (right) incubated with p-nitrophenyl butyrate (pNPB) (top row) and furimazine (fur.) (bottom row). (c) Titration of CATCH-cutinase (red circles, left y-axis) and CATCH-NL (green squares, right y-axis) activity into dual-enzyme gels (N=3) (d) Schematic of CATCH-U and CATCH-AdsA or CATCH-IDO dual-enzyme β-sheet fibrils. (e) Schedule to evaluate inflammation following subcutaneous MSU challenge in C57BL/6J mice pretreated with PBS (red circle), CATCH-U gel (green triangle), CATCH-AdsA gel (light blue inverted triangle), CATCH-IDO gel (orange inverted triangle), CATCH-U + AdsA gel (dark teal diamond), and CATCH-U + IDO gel (pink diamond). (f) Fold change in paw diameter 24 hours after MSU injection, normalized to day 0 measurement before MSU injection. N=6-11, ANOVA with Dunnett’s multiple comparisons to PBS (*) or CATCH-U gel (α). (g) Paw uric acid (solid symbols, left y-axis) and IL-1β (open symbols, right y-axis) in MSU-challenged paw. N=4-9, ANOVA with Dunnett’s multiple comparisons to PBS within each measurement. *P ≤ 0.05, **P ≤ 0.01 ***P ≤ 0.001, ****P≤ 0.0001. All data are presented as mean ± s.d.

In the MSU crystal-induced gout model, single-enzyme gel formulations with CATCH-U, CATCH-AdsA, or CATCH-IDO decreased initial paw swelling compared to PBS control (Figure 6d-f). Compared to CATCH-U gels, CATCH-AdsA gels were less effective while CATCH-IDO gels were more effective at reducing initial swelling. Dual-enzyme gels formulated with both CATCH-IDO and CATCH-U decreased initial paw swelling to the greatest extent, whereas CATCH-AdsA did not provide any additional benefit to CATCH-U (Figure 6f). Dual-enzyme gels of CATCH-U and CATCH-IDO more robustly decreased neutrophil degranulation 24 h after MSU challenge (Supplementary Fig. 11). The neutrophil inhibiting effect of the dual-enzyme gels paralleled their effectiveness for simultaneously dissolving MSU crystals and reducing IL-1β levels when compared to single enzyme CATCH- U or CATCH-IDO gels (Figure 6g).

## 3. Conclusion

This report demonstrates the successful realization of CATCH enzyme-peptide gels for localized therapeutic biocatalysis. CATCH-tagged fusion enzymes non-covalently integrate into β-sheet fibrils that are formed by CATCH(+/-) peptide pairs. At millimolar concentrations, these enzyme-bearing fibrils entangle into physically crosslinked gels^14^. Integration of five distinct CATCH fusion enzymes into functional single-enzyme or dual-enzyme gels showcased the modularity of this platform. By providing enzyme interchangeability and prolonged local retention, CATCH enzyme-peptide gels afford a generalizable vehicle that revitalizes a systemically-ineffective therapeutic enzyme with uricase, advances candidate immunomodulatory enzymes with AdsA and IDO, and provides multipronged therapy with dual-enzyme gels of uricase and IDO.

This work establishes supramolecular biocatalytic gels for in vivo use, expanding on a rich history of non-therapeutic biocatalytic gels for various industrial application such as chemical synthesis, food processing, biosensing, and environmental remediation.^12,51,52^ Prior to the last decade, enzyme immobilization was dominated by simple adsorption and covalent conjugation strategies that are often difficult to control and result in inconsistent active enzyme concentration. Non-covalent immobilization of enzymes within gels via supramolecular assembly can circumvent these challenges.^11^ The electrostatically controlled selective co-assembly of CATCH(+), CATCH(-), and CATCH fusion enzyme at physiologic pH, ionic strength, and temperature avoids denaturing stimuli, crosslinkers, or chemical modifications that can compromise enzyme catalytic activity. The triggered assembly mechanism results in titratable biocatalytic activity for both single- and dual-enzyme formulations by simply varying the fusion enzyme amount or type(s) during assembly. Supramolecular peptide gels, such as CATCH, are well positioned for in vivo applications because they are made of natural, biodegradable amino acids and are generally well tolerated.^14,53^ Finally, the shear thinning and recovery properties of CATCH gels,^14^ which are common among supramolecular peptide biomaterials,^54^ enable focal delivery through a fine gauge needle at the desired tissue site.

Direct local injection of a biodegradable enzyme-gel depot can replicate natural transient local enzyme expression better than systemic or targeted delivery approaches. Depots that prolong local enzyme retention can be achieved via fusion to anchoring domains with affinity for tissue ligands^46,55^ or by encapsulation in controlled release vehicles that leverage porosity or enzyme-carrier interactions for extended release.^56–63^ The CATCH gel system simplifies localized delivery by providing non-covalent enzyme immobilization in a carrier that is agnostic to tissue interactions and cargo properties. The 11 amino acid CATCH tag is amenable to proteins of disparate shapes, sizes, charges, and quaternary structures, as demonstrated with GFP (monomer, 26.6kDa, negative)^13^, cutinase (monomer, 22.6kDa, near neutral), NanoLuc^®^ (monomer, 19.1kDa, negative), IDO (monomer, 47.5 kDa, near neutral), AdsA (probable dimer, 87.4kDa, positive), and uricase (tetramer, 136.9 kDa, negative). Installation of the fusion enzyme directly into the β-sheet fibrils of the CATCH gel provided up to weeks of localized biocatalysis in vivo.

It is the localized deposits of MSU crystals in tissue that give rise to the inflammation and pain characteristic of gout flare, yet currently available therapies primarily act systemically to lower serum uric acid levels. So, although systemic administration of Pegloticase can be effective at reducing serum uric acid, it often fails to rapidly and completely dissolve MSU crystals^18^. Incomplete resolution of MSU crystals is thought to underlie recurrent flare, in addition to inducing systemic inflammation associated with the risk of developing comorbidities during inter-critical phases^64,65^. Although multiple bi-weekly infusions of Pegloticase over months can sufficiently reduce systemic serum uric acid levels to drive localized crystal deposits into solution^29^, many patients must discontinue treatment due to infusion reaction and increasing risk of anaphylaxis over many doses^18,32^. In contrast, localized delivery of uricase in the CATCH gel concentrates the therapeutic to the afflicted area, resulting in rapid dissolution of MSU crystals, as shown here with a single dose of CATCH-U gel in two mouse models of gout. ADAs raised against CATCH-U gels did not neutralize enzyme activity nor increase the risk of anaphylaxis, as they were skewed away from IgG1 and not IgE. Finally, as an alternative to infusion therapies, like Pegloticase^28^, which must typically be given at specialized infusion clinics over several hours, CATCH gels were designed to be administered in minutes via localized bolus injection through a fine-gauge needle, thereby greatly reducing risk of infection and need for specialized care. Overall, these data establish the CATCH gel as a broadly useful and easily-administered vehicle for localized delivery of one or more enzymes, enabling effective therapeutic biocatalysis in proximity to disease, while avoiding or minimizing adverse systemic side-effects.

## 4. Materials and Methods

### Recombinant protein expression and lysis

Previously published methods were adapted to generate experimental proteins^13,46,55^. All proteins were recombinantly expressed in an *E. coli* host system. Proteins include: CATCH-cutinase, CATCH-NL, WT NL, CATCH-IDO, WT IDO, CATCH-AdsA and CATCH-uricase. The fusion proteins consist of the enzyme covalently fused to the CATCH(-) peptide on the N terminus using a serine-glycine linker, and a 6-histidine tag on the C terminus for purification. The genes for these proteins were designed and ordered through GenScript in pET-21(+) vectors. Plasmid was transformed into One Shot^TM^ TOP10 Chemically Competent *E. coli* (ThermoFisher, C404003) and plated on 100 µg/mL ampicillin LB/agar plates overnight at 37 °C. Isolated colonies were selected and cultured in 5 mL of LB media with 100 µg/ mL ampicillin overnight in an orbital shaker at 225 rpm and 37 °C. Recombinant DNA was recovered with a Plasma Miniprep Kit (Qiagen) and sequenced using the Sanger method (GeneWiz). Positive DNA sequences were then transformed into Origami B (DE3) *E. coli* (Novagen, 70-837-4) and plated on ampicillin (100 µg/mL) and kanamycin (50 µg/mL) LB/agar plates overnight at 37 °C. Positive clones were picked and used to inoculate 5 mL of LB or 2xTy media with ampicillin (100 µg/mL) and kanamycin (50 µg/mL). The 5 mL of solution was cultured overnight at 37 °C on an orbital shaker (225 rpm) and then sub-cultured into 1 L of 2xTY media, consisting of 16 g tryptone, 10 g yeast extract, and 5 g NaCl, with ampicillin (100 µg/mL) and kanamycin (50 µg/mL) at 37 °C and in an orbital shaker until an optical density of 0.8 or greater at 600 nm was reached. To induce protein expression, the cultures were then supplemented with 0.5 mM isopropyl β-D-1-thiogalactopyranoside and incubated at 18 °C in an orbital shaker (225 rpm) for 18 h. Bacteria was then pelleted via centrifugation at 4 °C and 11,300 x g for 10 min using a Sorvall RC 6 Plus Superspeed Centrifuge. For CATCH-AdsA, bacteria pellet was resuspended in 20mL 50 mM HEPES and 100 mM NaCl, while the other protein-expressing bacteria was resuspended in 20mL 1x PBS. The bacteria were then centrifuged again using the same settings and the supernatant was discarded. CATCH-IDO and WT IDO cell pellets were mechanically resuspended in 1x PBS with Pierce™ protease inhibitor tablet (ThermoFisher) and lysed by sonication (15s on, 45s off, 10 min cycling time; ThermoFisher). All other cell pellets were mechanically resuspended and lysed in B-PER™ bacterial protein extraction reagent (ThermoFisher Scientific, 4 mL per gram of bacteria). Suspensions were incubated with lysozyme (50 mg/mL) and DNAse I (2400 units/mL) on a rocker for 20 minutes at room temperature. Lysates were then centrifuged at 42600 x g and 4°C for 15 minutes. The supernatants were collected by decanting for further purification.

### Cloning WT uricase

Template DNA was extracted from the CATCH-uricase TOP10 *E. coli* using a Plasma Miniprep Kit (Qiagen). WT uricase was cloned by deletion of the CATCH(-) tag using the Phusion^TM^ Site-Directed Mutagenesis Kit (Thermo Scientific), according to the manufacturer’s instructions. The new plasmid was transformed into Origami B (DE3) *E. coli*, expressed, and purified as described above and below.

### Purification of CATCH-AdsA

The His-tagged protein was purified by immobilized metal affinity chromatography (IMAC) using HisTrap FF Crude Prepacked Columns (Cytiva) connected to an ÄKTA Pure FPLC system (GE Healthcare). Fusion protein was eluted from the column using 20-250 mM imidazole gradient in buffer containing 50 mM HEPES and 100 mM NaCl. The imidazole in the solution was removed using Amicon Ultra Centrifugal Filters with a 10kDa cutoff (MilliporeSigma). IMAC and centrifugal filtration were repeated following the same conditions. The protein was purified through size exclusion chromatography (SEC) on the FPLC using a HiLoad^TM^ 26/600 Superdex 200 pg column. Using absorbance at 280nm, the protein was eluted and collected in 50 mM HEPES and 100 mM NaCl buffer.

### Purification of CATCH-cutinase, CATCH-NL, WT NL, CATCH-IDO, WT IDO, CATCH-uricase, and WT uricase

Proteins were purified following the same IMAC methods and centrifugal outlined above but with the imidazole gradient in 1x PBS. One round of IMAC was done for each protein. The imidazole solution was removed using 1x PBS and Amicon Centrifugal Filters with a 10kDa cutoff. Purification for all proteins was verified by SDS-PAGE gel, (30-50 min, 150-200 V) with Coomassie blue staining for 30-45 minutes, and destaining for 30-60 minutes.

### Endotoxin control

Endotoxin content of the recombinant proteins was reduced below 1.0 EU/mL using cloud point precipitation of Triton X-114 or Endotoxin Removal Solution (Sigma). For CATCH-AdsA, MnCl_2_ was added to the protein for a final concentration of 0.5 mM MnCl_2_ in 50 mM HEPES and 100 mM NaCl buffer. All other proteins were formulated in 1x PBS. Final endotoxin content was confirmed below 1.0 EU/mL, determined with Pierce™ LAL Chromogenic Endotoxin Quantitation Kit (ThermoFisher) or Chromo-LAL Endotoxin Quantification (Cape Cod Inc.), according to the manufacturer’s instructions.

### CATCH peptide stock solutions

A library of CATCH peptide variations exists based on the identity and number of the charged residues.^13,14,66^ This work focuses on: CATCH(+) [Ac- QQKFKFKFKQQ-Am] and CATCH(-) [Ac-EQEFEFEFEQE-Am]. The complementary peptides were synthesized and purified by GenScript (Supplementary Fig. 12 and 13). Lyophilized peptides were dissolved separately in endotoxin free water at a concentration of 24mM determined by weight. Sodium hydroxide was added to CATCH(-) to a pH ∼6.8 to aid dissolution. Both peptides were sonicated for 5 minutes to ensure complete dissolution. Phenylalanine absorbance (258nm) was measured by NanoDrop (ThermoFisher) to determine peptide concentration. For in vivo applications, endotoxin was confirmed below 1.0 EU/mL, determined with Pierce™ LAL Chromogenic Endotoxin Quantitation Kit (ThermoFisher) or Chromo-LAL Endotoxin Quantification (Cape Cod Inc.), according to the manufacturer’s instructions.

### Gel preparation

CATCH(+) was diluted with buffer to 12mM, while CATCH(-) was diluted with buffer and CATCH-fusion protein to 12mM. For CATCH-AdsA gels, 50 mM HEPES, 100 mM NaCl, 0.5 mM MnCl_2_ buffer was used, while 1x PBS was used for gel formation for all other proteins. Final concentrations of protein within the gel varied based on experiment. Gels were formulated to 12mM total peptide either by pipetting equal volumes of each stock solution on a surface (i.e. 96-well plate) or by drawing up equal volumes of each stock solution into the barrel of a syringe. In either case, the gels cured for 1 hour at room temperature before proceeding. For formation in the syringe, the 12mM peptide solutions were heated for 5 minutes before being sequentially drawn into the syringe to allow for more thorough mixing within the confines of the barrel. CATCH-AdsA was heated at 68°C, CATCH-IDO was heated at 45°C, and CATCH-NL, CATCH-cutinase, and CATCH-uricase were heated at 55°C. Allergy syringes with 27G or 30G needles (BD) were used.

### Animal housing conditions and strains

Animal protocols were approved by the Institutional Animal Care and Use Committee (IACUC) at the University of Florida and monitored by the Association for Assessment and Accreditation of Laboratory Animal Care (AAALAC). Procedures were performed following the Guidelines for Care and Use of Laboratory of the University of Florida and USDA guidelines. Mice were housed up to 5 in a cage, provided reverse osmosis water and standard diet, 14hr:10hr dark:light cycle, 68-79°F room temperature and 30-70% relative humidity. All mice were 8- to 15-weeks old at the start of experimentation. Female C57BL/6J mice (The Jackson Laboratory, #000664) were used. Breeding pairs of uricase knock out mice^23^ (UOx KO, B6;129S7-Uoxtm1Bay/J, The Jackson Laboratory #002223) were generously provided by Dr. Eric Gaucher from Georgia State University, and a colony was initiated. UOx KO mice were bred in-house and maintained on allopurinol (0.2 g/L, Patterson Vet Supply) *ad libitum* in their drinking water. Unless stated, all UOx KO animals were removed from allopurinol water at the beginning of experiment. Both female and male UOx KO mice were used for experiments. Mice were anesthetized with 2-3% isoflurane. The method of sacrifice was CO_2_ inhalation with cervical dislocation for secondary confirmation.

### CATCH-cutinase gel photos

100μL CATCH gels with and without 5 μM CATCH-cutinase (N=1) were formulated in 2mL glass vials and cured for 1 hour at room temperature. 20 μL of 2 mM pNPB in 1x PBS was added to each gel, the gels were inverted, and photos were taken with a digital camera (iPhone 16 Pro).

### CATCH-cutinase gel kinetics

10 μL gels with 5nM CATCH-cutinase were formed in 96-well plates. Then 100 μL of solution containing varying concentrations of pNPB (0-2.5mM) in 1x PBS was added to each well. Absorbance was read at 405nm for 60 minutes using a SpectraMax M3 plate reader (Molecular Devices). The slopes of the linear region of the kinetic measurements were determined and plotted against pNPB concentration. The Michaelis-Menten model in GraphPad was used to determine Michaelis constant, Km, and maximum velocity, Vmax.

### CATCH-cutinase gel activity titration

10 μL gels with varying concentrations of CATCH-cutinase (0-100nM) were formed in 96-well plates. Then 90 μL of 1mM pNPB in 1x PBS was added to each well. Absorbance was read at 405nm for 60 minutes. The slopes of the linear region of the kinetic measurements were determined and plotted against CATCH-cutinase concentration.

### Activity of CATCH-NL gels

NanoLuc^®^ is an engineered deep sea shrimp luciferase variant developed by Promega^67^. 400 μL CATCH gels with and without 100nM CATCH-NL were formulated in 2mL glass vials and cured for 1 hour at room temperature. 20 μL Nano-Glo furimazine (Promega) (diluted 2:5 with 1x PBS) was added to each gel, the gels were inverted, and photos with an iPhone 15 Pro were taken. Additionally, 10 μL CATCH gels with and without 100nM CATCH-NL were formed in triplicate in a white 96-well plate, cured for 1 hour, and washed 3 times with 200 μL 1x PBS. 90 μL 1x PBS was added to each gel for a total well volume of 100 µL. 50 μL Nano-Glo furimazine (diluted 1:50 with 1x PBS) was added to the wells and the luminescence was measured. Peak luminescence is reported.

### CATCH-NL retention in gel

CATCH-NL was immobilized within 10 μL gels to a final concentration of 5nM in a white 96-well plate and incubated for 24 hours. 90 μL of 1x PBS was added to each gel and incubated for another 24 hours. 80 μL of the supernatant was transferred to a separate well. An additional 80 μL 1x PBS was added to each gel, and 8 hours later 80 μL of supernatant was again removed and added to a separate well. Finally, 80 μL 1x PBS was again added to each gel and the luminescence of the gels, as well as the supernatants from 24- and 32-hour timepoints, were immediately measured after addition of 50 μL Nano-Glo furimazine (diluted 1:50 with 1x PBS).

### CATCH-NL in vivo imaging

Previously published methods were adapted to measure in vivo bioluminescence of CATCH-NL.^55^ While anesthetized, C57BL/6J mice received a single 20 μL subcutaneous injection of CATCH gels formulated with 50nM (N=4-6), 5nM (N=3), 0.5nM (N=4), or 0.05nM (N=3) CATCH-NL, or 50nM soluble CATCH-NL in 1x PBS (N=4), into the top of the hind paw. Five minutes later, 20 μL Nano-Glo furimazine (diluted 1:50 with 1x PBS) was injected subcutaneously into the same paw. Immediately after, mice were anatomically positioned with the hind paw facing the camera of an IVIS Spectrum In Vivo Imaging System (PerkinElmer) and bioluminescence images were taken. The Nano-Glo furimazine injections and IVIS imaging were repeated under anesthesia every 1-3 days over a 22-day period or until no signal above baseline was detected. Bioluminescent images were captured using an open emission filter, 1s exposure time, subject size 1cm, field-of-view B (6.6cm), medium binning (factor of 8) resolution, and a 1 F/stop aperture. Signal was quantified as photon flux (photons/s) using a round region of interest (ROI) in the Living Image analysis software (Revvity). The ROI captured the whole hind paw, stopping at the ankle, and the same ROI was used for each animal and each time point. Light intensities for each time point and animal were normalized to the same scale (min = 1.0e7 and max = 5e7) and set as pseudo color— where purple and red represented the least and most intense light respectively. To determine the average background signal (dotted line on graphs), naive contralateral paws of mice (N=8) received one subcutaneous injection of 20 μL Nano-Glo furimazine (diluted 1:50 with 1x PBS) and were imaged and analyzed as described above. Semi-log nonlinear regressions (Y=10^mX+b^) were calculated for longitudinal luminescence measurements of mice that received varying concentrations of CATCH-NL in gel formulation, where the inverse of the slope is used as the rate of clearance.

### CATCH-cutinase and CATCH-NL dual-enzyme gels

100 μL CATCH gels were formulated in 2mL vials containing the following: 5 μM CATCH-cutinase (N=1), 2.5 μM cutinase and 200nM CATCH-NL (N=1), and 400nM CATCH-NL (N=1). The gels cured for 1 hour at room temperature. 20 μL 2mM pNPB in 1x PBS was added to each gel, the gels were inverted, and photos were taken with a digital camera (iPhone 16 Pro). Then 2 μL furimazine (diluted 2:3 in 1x PBS) was added to each gel, the gels were inverted, and digital photos were taken again. In a separate study, CATCH gels (10 μL) were formed on a transparent 96-well plate in triplicate containing the following ratios of CATCH-cutinase to CATCH-NL: 4:0 (100nM:0nM), 3:1 (75nM:25nM), 2:2 (50nM:50nM), 1:3 (25nM:75nM), and 0:4 (0nM:100nM). The gels cured for one hour, then 90 μL pNPB in 1x PBS was added to each gel, for a final concentration of 1mM pNPB in each well. The reaction was monitored at an absorbance of 410 nm for one hour. Cutinase reaction velocity was calculated using the linear region of each reaction. Immediately after, 20 μL Nano-Glo furimazine (diluted 1:50 with 1x PBS) was added to the gels and luminescence from NL activity was measured (1000 ms integration time).

### CATCH-U gel activity with MSU crystals

MSU crystals were prepared as previously described^68^, with more detail in the supplement. 20 μL gels were formed with or without 16.5 μM CATCH-U in syringes. After the gels were cured at room temperature for an hour, they were injected into a 48-well plate and allowed to rest for 10 minutes at 4°C. The gels were washed three times with 200 μL 1x PBS. Then 200 μL 1x PBS with 0.625 g/L MSU and 2250 U/mL catalase was added to each well. The gels were imaged immediately using a Zeiss Axio Observer with the Ph3 condenser annulus. The gels were incubated for 24 hours at 37°C with gentle rocking and imaged again.

### CATCH-U gel activity with uric acid

20 μL gels were formulated with or without 16.5 μM CATCH-U in syringes. After the gels were cured at room temperature for an hour, they were injected into a UV-transparent 96-well plate (Corning) and allowed to rest for 15 minutes at 4°C. The gels were incubated with 200 μL 1x PBS for 1 hour at 37°C to remove any unincorporated peptide or enzyme. Then the PBS wash was removed, 200 μL 0.4mM uric acid was added to each well, and the absorbance of uric acid (293nm) was read immediately at 37°C for one hour. A linear regression of a uric acid standard curve was used to convert absorbance into uric acid concentration.

### MSU crystal inflammation paw caliper measurements

While anesthetized, C57BL/6J mice received a 20 μL injection subcutaneously into the top of the hind paw of one of the following treatments: 1x PBS (N = 9), 16.5 μM WT uricase in 1x PBS (N=8), empty CATCH gel (N=14), or 16.5 μM CATCH-U gel (1x, N=9). Two days later, the mice were challenged in the top of the same hind paw with 0.5mg MSU crystals suspended in 20 μL 1x PBS, a common model of gouty inflammation^19,20^. Paw diameter was measured with digital calipers prior to treatment and then everyday thereafter until the end of the study. Fold change in paw diameter was determined by normalizing each measurement to the measurement immediately before MSU crystal injection (t_x_/t_0_). Linear regression of each animals’ paw measurements were extrapolated to calculate time to baseline, when t_x_/t_0_ = 1. Regressions where R^2^ < 0.7 and outliers identified by ROUT analysis were removed. These methods were repeated with UOx KO mice maintained on *ad libitum* allopurinol water (0.2g/L) (N=6) or treated with 20 μL 1x PBS (N=11), empty CATCH gel (N = 12), or 33 μM CATCH-U gel (2x, N=13). Blood was collected from these mice by facial vein bleed at the beginning of the experiment and 5 days after MSU injection. Blood was collected from naive C57BL/6 mice (N=4) for comparison. Allopurinol was added to the blood (1 μL, 1.2mM) to inhibit endogenous xanthine oxidase^69^. Serum was separated and uric acid was quantified with Uric Acid Assay Kit (abcam) according to manufacturer guidelines. The paw caliper measurement methods were repeated again with C57BL/6J mice treated with 20 μL PBS (N=11), gel with 16.5 μM CATCH-U (N=6), gel with 18 μM CATCH- AdsA (N=6), gel with 16.5 μM CATCH-U and 18 μM CATCH-AdsA (N=6), gel with 15.8 μM CATCH-IDO (n=7), and gel with 16.5 μM CATCH-U and 15.8 μM CATCH-IDO (N=8).

### Paw IL-1β ELISA

While anesthetized, C57BL/6J mice received a 20 μL injection subcutaneously into the top of the hind paw of one of the following treatments: 1x PBS (N=4), empty CATCH gel (N=4), or 16.5 μM CATCH-U gel (N=4). Two days later, the mice were challenged with 0.5mg MSU crystals suspended in 20 μL 1x PBS. Treated paws were harvested 24 hours after MSU challenge and minced for homogenization in 1mL RIPA lysis buffer (ThermoFisher) with Pierce™ protease inhibitor tablet (ThermoFisher), 200mg garnet shards, and one 6mm zirconium bead using the Beadbug homogenizer (5 cycles, 30 second on, 30 seconds off) (Benchmark). Naive paws (N=4) were also processed for comparison. Homogenates were incubated on ice for 45 minutes and then centrifuged for 20 minutes at 12,000xg and 4°C. Supernatants were removed and protein content was quantified by Pierce™ BCA Protein Assay (ThermoFisher). Samples were normalized to 1000mg/mL protein and analyzed for IL-1β with ELISA (DuoSet, R&D Systems, Cat. No. DY401) according to manufacturer guidelines. The methods were repeated with C57BL/6J mice treated with 20 μL PBS (N=9), gel with 16.5 μM CATCH-U (N=4), gel with 15.8 μM CATCH-IDO (N=6), and gel with 16.5 μM CATCH-U and 15.8 μM CATCH-IDO (N=7). Outliers identified by Grubbs’ analysis were removed where appropriate.

### Paw MSU crystal burden

While anesthetized, UOx KO mice received a 20 μL injection subcutaneously into the top of the hind paw of one of the following treatments: 1x PBS (N=6), empty gel (N=4), 4.13 μM CATCH-U gel (0.25x, N=5), 8.25 μM CATCH-U gel (0.5x, N=5), 16.5 μM CATCH-U gel (1x, N=6), or 33 μM C-U gel (2x, N=4). Two days later, the mice were challenged subcutaneously in the hind paw with 0.5mg MSU crystals suspended in 20 μL 1x PBS. One hour after MSU injection, a cohort of mice received 33 μM soluble CATCH-U (2x, N=6). One day after MSU challenge, the paws were collected, weighed, and frozen at –80°C. Untreated contralateral paws (N=12) were also taken. Paws were thawed, diced into small pieces, and homogenized in 6mL water with the TissueRuptor (Qiagen) for 2 minutes at maximum speed and 1 minute at half-speed. A standard curve of MSU crystals (0.5mg, 0.25mg, 0.125mg, 0mg) were similarly homogenized. Tissue homogenates were centrifuged at 400xg for 5 minutes and supernatants were collected and passed through a 0.22µm filter. 100 μL of each sample or standard were plated in duplicate on a UV-transparent 96-well plate (Corning). The absorbance of uric acid was read at 293nm and linear regression of the standard curve was used to convert absorbance into uric acid concentration, which was then normalized to paw weight. Paw uric acid was also measured 5 days after MSU challenge in UOx KO mice treated with 20 μL 1x PBS (N=11), empty gel (N=12), 33 μM C-U gel (2x, N=12). MSU burden in all contralateral paws was also measured. The methods were repeated with C57BL/6J mice treated with 20 μL PBS (N=5), gel with 16.5 μM CATCH-U (N=6), gel with 15.8 μM CATCH-IDO (N=6), and gel with 16.5 μM CATCH-U and 15.8 μM CATCH-IDO (N=6). Outliers identified by Grubbs’ analysis were removed where appropriate.

### CATCH-U immunogenicity IgG titer

To measure the generation of antibodies against CATCH-U, C57BL/6 mice (N=5 per group) received a 100 μL subcutaneous scruff injection of 16.5 μM CATCH-U soluble in 1x PBS, in a CATCH gel, or emulsified in TiterMax^®^ Gold Adjuvant (Sigma-Aldrich). All mice received a 50 μL 16.5 μM CATCH-U scruff injection booster on day 28. Blood was drawn weekly from the facial vein through day 42. Sera from day 27 and 42 were analyzed for anti-uricase IgG titer by ELISA adapting established methods^11,14^. Briefly, plates were coated overnight at 4°C with 100 μL 1x PBS or CATCH-U (1 ug/mL) in 1x PBS. Plates were washed three times with 0.5% Tween-20 in PBS (PBST) and blocked with 1% bovine serum albumin (BSA) in PBST. Mouse sera were diluted with 1% BSA to 1:100, 1:1000, 1:10000, up to 1:10,000,000. Diluted serum (100uL) was added to blocked wells and incubated for 1 hour at room temperature. Serum was removed and plates were washed three times with PBST. Peroxidase-conjugated goat anti-mouse IgG (Abcam) was added to each well (100 uL, 1:5000 in PBS with 1% BSA) and then incubated for 1 hour at room temperature. Plates were washed five times with PBST. Plates were developed with 100 μL TMB substrate (ThermoFisher) for 30 minutes at room temperature. Stop solution (100 uL, 0.16 M sulfuric acid) was added to each well and absorbance was measured at 450 nm. To calculate the titer thresholds (TT), the PBS-coated control wells run alongside each individual serum were averaged for each titer, standard deviation calculated, and then TT defined as: TT = mean + 5*SD. Any serum absorbance readings greater than the TT for that titer were counted as positive and assigned the titer value (2, 3, 4, etc.).

### CATCH-U ADA isotyping

IgE and IgG isotyping was done on serum collected from immunogenicity mice on day 42 using Mouse IgE Uncoated ELISA Kit (ThermoFisher, 88-50460-22) and Mouse Monoclonal Antibody Isotyping (ThermoFisher, ISO2) respectively, according to manufacturer guidelines. For the IgE ELISA, plates were coated with the protein at 1 ug/mL overnight at 4°C. Serum were diluted to 1:25 as suggested by the manufacturer, and then the ELISA proceeded exactly as suggested, with a standard curve of IgE serving as our limit of detection for the assay. For the isotyping ELISA, plates were coated with protein overnight at 1 ug/mL (50 uL), then washed three times with PBST, then blocked for one hour with 100 μL of 1% acetylated BSA in PBST. The blocking solution was removed and then 50 μL of serum diluted in 1% acetylated BSA/PBS to 1:500 (except for the adjuvanted group, which was diluted to 1:5000) was added to the wells alongside PBS coated wells serving as control. After one hour, serum was removed and the wells were washed three times with PBST and 50 μL the detection antibodies were applied at a concentration of 1:1000 diluted in acyBSA/PBS for one hour. The wells were washed with PBST three times, then 50 μL of the secondary antibody, Rabbit Anti-Goat IgG Antibody, HRP conjugate (ThermoFisher, AP106P) was added at a concentration of 1:5000 diluted in the 1% acetylated BSA/PBS solution for 15 minutes. The wells were washed three times and then developed for 5 minutes with 50 μL of TMB solution, followed by 50 μL of the stop solution, 0.16 M sulfuric acid, and then reading at 450 nm. PBS control wells were subtracted from the test wells. Each IgG isotype was normalized to the measured IgM for animal.

### ADA neutralization of uricase activity

In a UV-transparent 96-well plate (Corning) soluble CATCH-U (20uL, 16.5uM) was incubated with PBS (20uL) or serum collected from immunogenicity mice on day 35 (20uL, diluted 1:10 in PBS) on a rocker for one hour at room temperature. Sera included TiterMax (N=5), CATCH-U gel (N=5), or soluble CATCH-U (N=5). For a positive control of uricase activity inhibition, oxonic acid (20uL, 8mM) was added to soluble CATCH-U (20uL, 16.5uM). Uric acid was added to each well (200uL, 0.4mM) and uric acid depletion was measured at 293nm at 37°C for 30 minutes. The linear region of each reaction was used to calculate reaction velocity, and a standard curve of uric acid was used to convert this absorbance into uric acid concentration.

### Soluble CATCH-AdsA activity

Soluble CATCH-AdsA protein kinetics was assessed using malachite green assay kit (Sigma) to measure free phosphate production in a 96-well plate. To analyze the substrate-dependent activity, 2 µM AdsA was incubated with ATP and ADP at 50 µM – 8 mM. After 5 minutes of incubation, the working reagent was added, and 5 min later absorbance was measured at 620 nm. Absorbance measurements were repeated every five minutes to measure the levels of free phosphate in the well over 30 minutes. The velocity of each reaction was measured as the slope of the linear region. Using the Michaelis-Menten model in GraphPad Prism, the Michaelis constant (KM) and maximum velocity (Vmax) were determined. Soluble CATCH-AdsA activity was further evaluated at various enzyme concentrations to determine the relative ATP turnover using the ATP Bioluminescence Assay Kit CLS II (Roche). A range of CATCH-AdsA concentrations (0, 1, 2, 9, and 20 uM) were incubated in a white 96-well plate with 10 μM ATP for two hours at room temperature. A standard curve of ATP was also placed on the plate at the start of the run. At the end of the two hours, luminescence was measured using the kit for ATP, where concentration of ATP in the test wells was determined using the standard curve.

### CATCH-AdsA gel activity

The ability of CATCH-AdsA gels to dephosphorylate ATP was tested by measuring ATP concentration bioluminescence as above. Empty and 9 μM CATCH-AdsA gels were prepared at 40 μL in syringes as described previously. Gels were injected into a white 96-well plate and incubated with 40 μL 2000 µM ATP in HEPES buffer for one hour. Approximately 15 minutes before the end of the hour, fresh luciferase reagent was prepared following the manufacturer’s instructions. Luciferase was added at 80 μL/well, and luminescence readings were taken, with an integration time of 1000 ms. The activity of CATCH-AdsA gels to dephosphorylate ATP was also assessed by measuring free phosphate generation using the malachite green assay kit (Sigma). Gels were formed with 2 μM CATCH-AdsA in a 96-well plate and incubated with 30 μM ATP for 1 hour. 80 μL 30 μM ATP alone was incubated for 1 hour as a control. 20 μL of the working reagent was added to each well. After 15 minutes, absorbance was measured at 620nm and compared to a standard curve.

### CATCH-AdsA gels and THP-1 cells

THP-1 cells (ATCC, TIB-202) were seeded into a 6-well plate at a density of 1 million cells per well and incubated at 37°C and 5% CO_2_. Cells media contained RPMI-1640 (Gibco), 10% FBS (Cytiva), 1% penicillin streptomycin (Gibco), 1x GlutaMax L-glutamine (Gibco), 1mM sodium pyruvate (Gibco), 10mM HEPES (Gibco), and 0.05 μM 2-mercaptoethanol. Cells were treated with 50 ng/mL phorobol-12-myristate-13-acetate (PMA) for 24 hours. Media was replaced and cells were incubated for another 24 hours. Cells were harvested and reseeded into a 96-well plate at a density of 300,000 cells per well. Cells were cultured overnight before being stimulated with 10 µg/mL lipopolysaccharide (LPS) for 18 hours. Culture media was replaced, and the cells were treated with 100 μL of the following: 1x PBS, 2mM ATP, 2mM ATP reacted for 2 hours with empty gels, or 2mM ATP reacted for 2 hours with 2 μM CATCH-AdsA gels. Cultures were incubated for 1 hour before the media was collected for analysis. IL-1β production was quantified using a human IL-1β ELISA kit (BD OptEIA) according to manufacturer’s protocol.

### CATCH-AdsA gels in LPS model immunohistochemistry

Cohorts of C57BL/6J mice received a 40 μL subcutaneous injection into the top of the hind paw of either empty gel (N=3) or 9 μM CATCH-AdsA gel (N=3), and 24 hours later 40 μL LPS (1 mg/mL) was injected into the same paw. Paws were harvested 24 hours after LPS injection. Neutrophil infiltration into the subcutaneous space of the hind paw was assessed with immunohistochemistry (IHC). The whole paws were fixed with 10% neutral buffered formalin, decalcified, cut bilaterally, and embedded in paraffin. The blocks were sectioned onto slides and stained with anti-neutrophil Ly6-B.2 antibody (Abcam, ab53457). Histological images were analyzed by a blinded individual. The observer created four rectangular regions of interest 400 μm long equally spaced along the length of the paw. The number of Ly6-B.2+ stained cells were manually counted in each region and normalized by the area. The average across the four sections was calculated and plotted for each animal.

### CATCH-AdsA gels in LPS model luminol measurements

Luminol was used to assess MPO activity from neutrophil infiltration into the injection site.^70^ Cohorts of C57BL/6J mice received a 40 μL subcutaneous injection into the top of the hind paw of either empty gel (N=10), gel with 2 μM CATCH-AdsA (N=9), gel with 5 μM CATCH-AdsA (N=2), or gel with 9 μM CATCH-AdsA (N=13), 24 h before 40 µL of LPS (1 mg/mL) paw challenge at day 0. On day 1, the animals received a 100 μL intraperitoneal (i.p.) injection of luminol solution (20.2 mg/mL in 0.9% NaCl saline). 10 minutes after, the mice were individually anatomically positioned with the hind paw facing the camera of the IVIS and bioluminescence images were taken. The luminol injection and imaging was repeated every 24 hours for a total of 4 consecutive days. Bioluminescent images were captured using an open emission filter, 1 minute exposure time, subject size 1cm, field-of-view B (6.6cm), large binning resolution, and a 1 F/stop aperture. Luminol emission signal was quantified as total flux using a round region of interest (ROI) using the Living Image analysis software. The ROI used captured the whole hind paw, stopping at the ankle. The same ROI was used for each animal and each time point.

### CATCH-IDO gel biochemical activity

Previously published methods were adapted to measure the catalytic activity of CATCH-IDO gels^45,71^. In a 96-well plate, 20 μL gels were formulated in triplicate with and without 10 μM CATCH-IDO. The gels were washed with 200 μL 1x PBS on a rocker for 10 minutes and the wash was removed. Then 30 μL 1x PBS and 50 μL master mix containing 0.36mM tryptophan, 4900 U/mL catalase, and the necessary electron donors, 18.2 μM methylene blue and 43.6mM ascorbic acid, was added to each well. N-formyl-kynurenine (NFK), the immediate precursor to Kyn, was measured at 321nm at 37°C for 20 minutes. Data was normalized to empty gel control.

### CATCH-IDO gels with LPS-stimulated dendritic cells

Methods to isolate mouse bone marrow derived dendritic cells and culture them with IDO were adapted from previously published methods^45^. Primary bone marrow cells were harvested from C57BL/6J mice and differentiated into dendritic cells with granulocyte-macrophage colony-stimulating factor (GMCSF) (20ng/mL, Biolegend) over 10 days. Cells were cultured in media containing DMEM/F12 with L-glutamine (Gibco), 10% FBS (Cytiva), 20ng/mL GMCSF, 1mM sodium pyruvate (Gibco), 10mM HEPES (Gibco), 1x NEAA (Gibco), 1% penicillin streptomycin (Gibco), and 50μM 2-mercaptoethanol. Dendritic cell viability and purity (CD11b-, CD11c+) were confirmed to be above 80% with flow cytometry prior to experimentation (Cytek Aurora, fixable viability dye eFluor 780 [APC-Cy7, eBiosciences], CD11b [M1/70, BV510, Biolegend], CD11c [N4180, BV711, Biolegend], MHCII [M5/114.15.2, BV421, Biolegend], CD80 [16-10A1, APC, eBiosciences], CD86 [GL1, PE-Cy7, eBiosciences]). Cells were seeded in 1mL media at 0.5x10^6^ cell/well in 12-well plates and treated with either 30 μL PBS, soluble CATCH-IDO (20.2uM), empty gels, or CATCH-IDO (20.2uM) gels for 24 hours and then stimulated with LPS (1 ug/mL) or 1x PBS overnight. Cytokine release (IL-12 p70 BD Biosciences and IL-6 DuoSet R&D Systems) was measured from the supernatant by ELISA. Cells treated with empty gels or CATCH-IDO gels without LPS stimulation had negligible levels IL-12 and IL-6 (data not shown).

### Anti-IDO immunofluorescence

Hair was removed from the backs of anesthetized C57BL/6J mice with clippers and depilatory cream 24 hours prior to injection. CATCH(+) peptide was labeled with 1% AlexaFluor750 NHS ester (ThermoFisher) for 30 minutes at room temperature. 20 μL gels were formed in syringes with a final concentration of 0.5% AlexaFluor750 without enzyme (N=1) or with 20 μM CATCH-IDO (N=1) and subcutaneously injected into the shaved backs of mice. The injection sites were harvested 2 hours later, embedded in O.C.T. Compound (Fisher), frozen in liquid nitrogen, and kept at -80°C. The tissue was cryo-sectioned to 15 μm thickness and stained with anti-human IDO antibody (Cell Signaling Technologies, D5J4E, #86630) and DAPI.

### Kynurenine ELISA

C57BL/6J mice were injected in the ear and subcutaneous space on the top of the hind paw with 20 μL empty gel (N=3-5) or CATCH-IDO gels (20 μM, N=2-3). One hour later the tissues were harvested, weighed, and frozen at -80°C. Naive tissues (N=2-3) were also harvested for comparison. Tissues were thawed, minced, and homogenized in 0.8mL methanol with 200mg garnet shards and one 6mm zirconium bead using the Beadbug homogenizer (5 cycles, 30 second on, 30 seconds off) (Benchmark). The homogenates were incubated on ice for 15-30 minutes and then centrifuged at 10,000-15,000xg for 10-15 minutes at 4°C. The supernatants were passed through a 0.22 µm filter and then the solvent was evaporated off. The isolate was reconstituted in 30 μL water and kynurenine was measured by ELISA (Immunsmol) according to manufacturer guidelines. Kynurenine levels were normalized to tissue weight.

### CATCH-IDO gels in IMQ-induced psoriasis

Previously published methods were adapted to establish the IMQ-induced psoriasis model^49,72^. Hair was removed from the backs of anesthetized C57BL/6J mice with clippers and depilatory cream. Each day for 10 days total, 5% IMQ cream (62.5 mg, Patterson Veterinary Supply) was applied to the backs of the mice. On the 3rd day of IMQ application, mice were treated with 5 subcutaneous injections spaced across the back of 20 μL PBS, CATCH-IDO gels (20uM), or empty gels (N=6 per group). Disease severity was measured each day using a modified version of the Psoriasis Area and Severity Index (PASI) where area of effect is not considered. Erythema (redness), scaling, and thickening were scored independently and on a scale of 0 to 4: 0, none, 1: slight, 2: moderate, 3: marked, 4: very marked. The cumulative score was reported as a measure of the severity of inflammation (scale 0-12). Digital images were taken of mouse back on day 7. An additional cohort of mice (N=8-12 per group) were euthanized on day 7 and the back skin and draining inguinal lymph nodes were collected for flow cytometry analysis.

### IMQ dermis flow cytometry

Cells from the back skin were isolated following a modified protocol from Lou et al.^73^ Three 1cm^2^ skin squares were harvested and then quartered and digested in Dispase II (5 mg/mL, Sigma) for 1 hour at 37°C. Dermis and epidermis were separated, and dermis layers were digested in collagenase P (1 mg/mL, Sigma) and DNAse I (1 mg/mL, Sigma) for 80 minutes at 37°C. Following digestion, cells from the dermis were passed through a 40µm cell strainer, counted, and pretreated with TruStain FcX (anti-mouse CD16/32, Biolegend). The cells were stained for stained for viability [fixable viability dye eFluor 780, APC-Cy7, eBiosciences], CD45.2 [104, AF700, Biolegend], CD11b [M1/70, BV510, Biolegend], CD11c [N4180, BV711, Biolegend], Gr-1 [RB6-8C5, BUV395, eBiosciences], Ly-6G [1A8-Ly6g, eF450, eBiosciences], TCRβ [H57-597, eF450, eBiosciences], TCRγδ [GL3, PE, Biolegend], CD4 [RM4-5, SuperBright 780, eBiosciences], CD8α [53-6.7, BV605, eBiosciences], CD44 [IM7, BV650, Biolegend], CCR6 [29-2L17, PE/Fire 810, Biolegend], and FoxP3 [MF23, PE-cf594, BD Biosciences]. Flow cytometry was performed with a Cytek 5-laser Aurora Cytometer and analysis was preformed performed using FlowJo (BD Biosciences). Gating strategy is outlined in Supplemental Information.

### Dual-enzyme gels in gout model luminol measurements

Cohorts of C57BL/6J mice received a 20 μL subcutaneous injection into the top of the hind paw of either 1x PBS (N=11), gel with 16.5 μM CATCH-U (N=6), gel with 18 μM CATCH-AdsA (N=6), gel with 16.5 μM CATCH-U and 18 μM CATCH-AdsA (N=6), gel with 15.8 μM CATCH-IDO (N=7), and gel with 16.5 μM CATCH-U and 15.8 μM CATCH-IDO (N=8). Two days later, the mice were challenged with hind paw injection of 0.5mg MSU crystals suspended in 20 μL 1x PBS. One day after MSU challenge, and only for this timepoint, the animals were assessed for MPO activity with luminol as described above.

### Statistical Analysis

Statistical analyses were performed using GraphPad Prism. Study-specific analyses are reported in figure captions using the following: non-significant (ns) P > 0.05, *P ≤ 0.05, **P ≤ 0.01, ***P ≤ 0.001, and ****P ≤ 0.0001. All data are presented as mean ± s.d. All t-tests reported are two-tailed.

## Acknowledgements

This research was supported by the National Institute of Health. Support to Gregory Hudalla is acknowledged from 5 R21 EB024762, NSF RAISE CBET-1743432, R01 DK129690. We would like to thank Dr. Jorge Mojica Santiago (Dept of Biomedical Engineering, University of Florida) for access to the rheometer and Animal Care Services and Veterinary Staff for supporting the animal studies. We would like to thank Dr. Eric Gaucher from Georgia State University for his generous donation of UOx KO breeder mice for colony establishment.

Histology was performed at the Molecular Pathology Core in the Department of Pathology, Immunology, and Laboratory Medicine at the University of Florida College of Medicine, RRID: SCR_016601. This work utilized an Olympus VS200 whole slide scanner purchased with an NIH-shared instrumentation grant S10OD032236.

Received: ((will be filled in by the editorial staff))

Revised: ((will be filled in by the editorial staff))

Published online: ((will be filled in by the editorial staff))

