## Supplementary Information for "Supramolecular Enzyme-Peptide Gels for Localized Therapeutic Biocatalysis"

#### Supplementary Methods

*Analytical size exclusion chromatography.* 500 μL solution of purified protein (CATCH-uricase, CATCH-AdsA, CATCH-IDO and WT IDO) was loaded onto Superdex Increase 75 or 200 10/300 GL column (Cytiva) connected to an ÄKTA Pure FPLC system. Using absorbance of 280nm and protein standard markers, the fusion protein molecular weight was determined, or elution volume was reported.

O*scillating Rheology*. Samples of CATCH gels were prepared in syringes as described above. Samples included empty gels (no enzyme), gels with 100nM CATCH-cutinase, and gels with 16.5 μM CATCH-uricase. Gel rheological properties were analyzed using an Anton Paar rheometer (MCR 302). The gels were injected onto the instrument's rough-coated plate, and the 8mm rough-coated parallel plate was lowered to the gel at 0.5mm. The linear viscoelastic region (LVR) was determined at 37°C using amplitude sweeps. Frequency sweeps (10-0.1 rad/s) were run in the LVR at 0.3% strain to determine the storage modulus (G') and loss modulus (G''). Damping factor was calculated as G'/G'' at angular frequencies 0.1, 1.0, and 10 rad/s.

*Effect of CATCH-tag on NL activity*. The activity of CATCH-NL and WT NL were compared at concentrations of 0-10nM. 50 μL of protein at each concentration was added to separate wells of white 96-well plate. Then 50 μL Nano-Glo furimazine (Promega) (diluted 1:50 with 1x PBS) was added and the luminescence was immediately measured using the SpectraMax M3 plate reader (Molecular Devices).

*Preparation of monosodium urate (MSU) crystals*. MSU crystals were prepared as previously described by Alberts et al[1]. 20mM uric acid (Alfa Aesar) was dissolved in boiling water containing 20mM NaOH. The solution was cooled to 60°C, adjusted to pH 9 with NaOH, and sterile filtered. The solution was left under mild agitation at room temperature for 24-72 hours to allow crystal formation. The crystals were then harvested on a sterile filter, washed with 70% ethanol, and dried overnight at 60°C. The crystals were stored dry at 4°C or resuspended in 1x PBS and stored at -20°C. Solutions of crystals were sonicated for 3-5 minutes to reduce crystal size to allow injection through a 25G-27G needle. Crystals were analyzed by microscopy to ensure correct formation of their needle-like structure. Endotoxin content was confirmed to be below 1.0 EU/mL, determined with Chromo-LAL Endotoxin Quantification (Cape Cod Inc.), according to the manufacturer's instructions.

*Temperature sensitivity of CATCH-uricase activity*. The ability of the pre-heated uricase to deplete uric acid was assessed by measuring the absorbance of uric acid at 293nm over time. CATCH-U at 16.5 μM was incubated for 5 minutes at room temperature, 37°C, 45°C, 50°C, 55°C, 60°C, 65°C, and 70°C. 20 μL pre-heated CATCH-U was added in triplicate to a UV-transparent 96-well plate with 200 μL of 0.4mM uric acid and absorbance was read for 30 minutes. To get relative activities, the reaction velocity was determined for each condition and normalized to reaction velocity at RT incubation.

*Effect of CATCH-tag on uricase activity*. The activity of CATCH-U was compared to the activity of WT uricase. 5 μL solution containing 16.5 μM of each protein was pipetted into a UV-transparent 96-well plate. Then 75 μL of uric acid at various concentrations (250-1000uM) was added to each well and absorbance was immediately read at 293nm for 10 minutes at 37°C. The slopes of the linear region of the kinetic measurements were determined and converted to reaction velocity (uric acid depletion per minute) using the slope of a uric acid standard curve. These activities were then plotted against initial uric acid concentration and the slopes of the CATCH-uricase and WT uricase were compared in GraphPad Prism.

*CATCH-uricase retention in CATCH gels*. 20 μL gels were formulated with 16.5 μM CATCH-U or with 16.5 μM WT uricase in syringes. The gels were injected into a 96-well plate, 200 μL 1x PBS was added to each well, and the plate was left at room temperature. A standard curve of CATCH-U and WT protein (200 μL/well) was also plated. 48 hours later 175 μL of the PBS supernatant was removed from each gel and plated in a black 96-well plate, along with 175 μL of each standard. The remaining 25 μL 1x PBS supernatant on the gels was discarded. 200 μL of fresh 1x PBS was added to each gel and rapidly pipetted to dislodge and homogenize the gel. 175 μL of the gel homogenate was plated in the black 96-well plate. Then tryptophan fluorescence (ex/em 280/345) was measured to quantify the protein content. A linear regression of the standard curves was used to convert fluorescence into protein content. Then percentages of each fraction, inside the gel vs in the supernatant, were calculated using the total quantified protein signal of the sample; i.e. percentage in the supernatant equals signal from supernatant divided by signal from supernatant plus signal from gel.

*CATCH-U-cy5.5 retention in vivo imaging*. CATCH-U and wild type uricase was fluorescently labeled with cyanine-5.5 (cy5.5) through maleimide-cysteine chemistry. The reactions were carried out with a 20x molar excess sulfo-cy5.5 maleimide (Lumiprobe) at 4°C overnight according to the manufacturing guidelines. Unreacted dye was removed by buffer exchange and filter centrifugation. Cohorts of mice were injected with 20 μL into the subcutaneous space on the top of the hind paw with either 16.5 μM WT uricase-Cy5.5 in 1x PBS (N=4) or CATCH gels with 16.5 μM CATCH-U-cy5.5 (N=5). The mice were individually anatomically positioned with the hind paw facing the camera of the IVIS and fluorescent images were taken immediately after injection. Images were taken again 30 minutes, 1 hour, and 3 hours later, then once per day for a 10-day period. Fluorescent images were captured using ex/em 675/720nm, 1 second exposure time, subject size 1cm, field-of-view B (6.6cm), small binning resolution, and a 2 F/stop aperture. Cy5.5

signal was quantified as radiant efficiency using a round ROI in the Living Image analysis software. The ROI used captured the whole hind paw, stopping at the ankle. The same ROI was used for each animal and each time point. Half-life was calculated in GraphPad prism.

*MSU-induced inflammation paw histology*. Paw tissue was harvested from a cohort of animals 144 hours after MSU crystal injection. These animals were treated with 20 µL of either gels with 16.5 µM CATCH-U (N=5), empty gels (N=4), or PBS (N=3) 48 hours prior to MSU injection. Cell infiltration into the paw was assessed with histology. The whole paws were fixed with 10% formalin and decalcified then cut bilaterally and embedded in paraffin. The blocks were sectioned and stained with hematoxylin and eosin (H&E). Histology images were analyzed by a blinded individual. The observer measured the thickness of immune infiltrate in the subcutaneous space of the hind paw. Measurements were taken at 0.4mm intervals across the length of the paw and averaged per animal. One contralateral, naïve paw was used as a baseline control, shown as a dashed line on the graph. Another group of mice were treated with 20 µL of either 16.5 µM CATCH-U gel, empty gel, or PBS (N=1 per group) 48 hours prior to MSU injection, and paw skin was harvested from animals 24 hours after MSU crystal challenge. Fresh skin tissue was embedded in O.C.T. Compound (Fisher), flash frozen in liquid nitrogen and stored at -80°C. The tissue was cryo-sectioned to 15 µm thickness and stained with H&E to assess MSU crystal burden and cell infiltration.

*Temperature sensitivity of CATCH-AdsA.* The activity of the pre-heated AdsA to dephosphorylate ATP was assessed by measuring free phosphate generation using the malachite green assay kit (Sigma) in a 96-well plate. CATCH-AdsA at 2 µM was heated for 5 minutes at room temperature, 37 °C, 50°C, 70°C, or 90°C. 80 µL of the pre-heated CATCH-AdsA and 30 µM ATP was incubated in triplicate for one hour. 20 µL of the working reagent from the kit was added. After 15 minutes, absorbance was measured at 620nm. Relative activity was calculated as the amount of free phosphates generated normalized to the amount of free phosphates generated at RT.

*pH sensitivity of CATCH-AdsA*. The activity of CATCH-AdsA gels to dephosphorylate ATP at varying pHs was assessed by measuring free phosphate generation using the malachite green assay kit (Sigma) in a 96-well plate. Sodium hydroxide was added to CATCH-AdsA to a final pH of 5, 5.5-6.0, 6.4, 7.0, and 7.6. Then 80 µL of each CATCH-AdsA was added to 30 µM ATP. After a 1-hour incubation, 20 µL of the working reagent from the malachite green assay kit was added, and 15 minutes later, absorbance was measured at 620nm

*CATCH-IDO vs WT IDO activity.* 50 µL soluble CATCH-IDO or WT IDO at 0.358 µM were plated in triplicate in a 96-well plate. Then 50 µL master mix containing 0.36mM tryptophan, 4900 U/mL catalase, 18.2 µM methylene blue, and 43.6mM ascorbic acid was added to each well. N-formyl-kynurenine (NFK) production was measured at 321nm at 37°C for 10 minutes. Absorbance was blanked to PBS buffer control.

*CATCH-IDO Temperature sensitivity*. Soluble CATCH-IDO at 0.358 µM was heated for 5 minutes at RT, 37 °C, 45°C, 50 °C, 55°C, and 60°C. In a 96-well plate, 50 µL of each pre-heated CATCH-

IDO sample was plated in triplicate. Then 50 μL master mix containing 0.36mM tryptophan, 4900 U/mL catalase, 18.2 μM methylene blue, and 43.6mM ascorbic acid was added to each well. NFK was measured at 321nm at 37°C for 20 minutes. Absorbance was blanked to PBS buffer control. Relative activity was calculated as the reaction velocity normalized to the reaction velocity at RT.

*IMQ lymph node flow cytometry.* Cells from the inguinal lymph nodes were isolated following a modified protocol from Hornsteiner et al[2]. Lymph nodes were dissected and incubated with collagenase P (250ug/mL, Sigma) and DNAse I (300ug/mL, Sigma) to obtain a single cell solution. Following digestion, cells were passed through a 40μm cell strainer, counted, and pretreated with TruStain FcX (anti-mouse CD16/32, Biolegend). The cells were stained for stained for viability [fixable viability dye eFluor 780, APC-Cy7, eBiosciences], CD45.2 [104, AF700, Biolegend], CD11b [M1/70, BV510, Biolegend], CD11c [N4180, BV711, Biolegend], Gr-1 [RB6-8C5, BUV395, eBiosciences], Ly-6G [1A8-Ly6g, eF450, eBiosciences], TCRβ [H57-597, eF450, eBiosciences], TCRγδ [GL3, PE, Biolegend], CD4 [RM4-5, SuperBright 780, eBiosciences], CD8α [53-6.7, BV605, eBiosciences], CD44 [IM7, BV650, Biolegend], CCR6 [29-2L17, PE/Fire 810, Biolegend], and FoxP3 [MF23, PE-cf594, BD Biosciences]. Flow cytometry was performed on a Cytek 5-laser Aurora Cytometer and analysis was preformed performed using FlowJo (BD Biosciences).

*Flow cytometry gating.* FSC/SSC was used to select the cell population and remove debris. The immune cell population was identified by single cells, live cells (fixable viability dye negative), CD45+ cells. Neutrophils were identified by CD11b+, TCRγδ-, GR-1+, and Ly-6G+. Frequency of neutrophils was reported as percentage of CD11b+, with one statistical outlier removed in CATCH-IDO group (final N=7) and no outliers removed from PBS group (N=10). Lymphoid cells were identified as CD11b- CD11c-. γδ T cells were identified as TCRγδ+. Activated and Th17-like γδ T cells were identified as CD44+ and CCR6+, respectively, and reported as a percent of γδ T cells (N=12 PBS, N=12 CATCH-IDO). CD8 T cells were identified as TCRβ+ and CD8+ and were reported as a percent of TCRβ+ cells, with one statistical outlier removed in the CATCH-IDO group (final N=11) and no outliers removed from PBS group (N=12). Helper T cells were identified as TCRβ+ and CD4+, and subtypes were reported as percent of CD4+ cells. Th17 cells were identified as CCR6+, with one statistical outlier removed in CATCH-IDO group (final N=11) and no outliers removed from PBS group (N=12). T regulatory cells were identified as Foxp3+. For the dermis Treg cell population, one statistical outlier was removed from PBS group (final N=7) and no outliers were removed from CATCH-IDO group (N=8). No outliers were removed from the Treg population in the dermis (N=12 PBS, N=12 CATCH-IDO). Outliers were identified by ROUT analysis.

*CATCH-Cutinase amino acid sequence*: cloned from *Fusarium solani* variant of cutinase

MAEQEFEFEFEQEGSGGGSGGSGGGGSGGSGEFMRSGLPTSNPAQELEARQLGRTTRDD
LINGNSASCADVIFIYARGSTETGNLGTLGPSIASNLESAFGKDGVWIQGVGGAYRATLG
DNALPRGTSSAAIREMLGLFQQANTKCPDATLIAGGYSQGAALAAASIEDLDSAIRDKIA
GTVLFGYTKNLQNRGRIPNYPADRTKVFCNTGDLVCTGSLIVAAPHLAYGPDARGPAPE
FLIEKVRAVRGSALEHHHHHH

*CATCH-NanoLuc amino acid sequence*: cloned from deep-sea shrimp luciferase variant developed by Promega

MAEQEFEFEFEQEGSGGGSGGSGGGGSGGSGEFMAVFTLEDFVGDWRQTAGYNLDQV
LEQGGVSSLFQNLGVSVTPIQRIVLSGENGLKIDIHVIIPYEGLSGDQMGQIEKIFKVVYPV
DDHHFKVILHYGTLVIDGVTPNMIDYFGRPYEGIAVFDGKKITVTGTLWNGNKIIDERLI
NPDGSLLFRVTINGVTGWRLCERILALEHHHHHH

*CATCH-Uricase amino acid sequence*: cloned from *Aspergillus flavus* variant of uricase

MAEQEFEFEFEQEGSGGGSGGSGGGGSGGSGEFMSAVKAARYGKDNVRVYKVHKDEK
TGVQTVYEMTVCVLLEGEIETSYTKADNSVIVATDSIKNTIYITAKQNPVTPPELFGSILG
THFIEKYNHIHAAHVNIVCHRWTRMDIDGKPHPHSFIRDSEEKRNVQVDVVEGKGIDIKS
SLSGLTVLKSTNSQFWGFLRDEYTTLKETWDRILSTDVDATWQWKNFSGLQEVRSHVP
KFDATWATAREVTLKTFAEDNSASVQATMYKMAEQILARQQLIETVEYSLPNKHYFEID
LSWHKGLQNTGKNAEVFAPQSDPNGLIKCTVGRSSLKSKLLEHHHHHH

*CATCH-AdsA amino acid sequence*: cloned from *Staphylococcus aureus* variant of AdsA

MAEQEFEFEFEQEGSGGGSGGSGGGGSGGSGGGQHTPMKAHAVTTIDKATTDKQQVPP
TKEAAHHSGKEAATNVSASAQGTADDTNSKVTSNAPSNKPSTVVSTKVNETRDVDTQQ
ASTQKPTHTATFKLSNAKTASLSPRMFAANAPQTTTHKILHTNDIHGRLAEEKGRVIGM
AKLKTVKEQEKPDLMLDAGDAFQGLPLSNQSKGEEMAKAMNAVGYDAMAVGNHEFD
FGYDQLKKLEGMLDFPMLSTNVYKDGKRAFKPSTIVTKNGIRYGIIGVTTPETKTKTRPE
GIKGVEFRDPLQSVTAEMMRIYKDVDTFVVISHLGIDPSTQETWRGDYLVKQLSQNPQL
KKRITVIDGHSHTVLQNGQIYNNDALAQTGTALANIGKITFNYRNGEVSNIKPSLINVKD
VENVTPNKALAEQINQADQTFRAGGGGSGGGGSHHHHHH

*CATCH-IDO amino acid sequence*: cloned from human variant of IDO

MAEQEFEFEFEQEGSGGGSGGSGGGGSGGSGEFMAHAMENSWTISKEYHIDEEVGFALP
NPQENLPDFYNDWMFIAKHLPDLIESGQLRERVEKLNMLSIDHLTDHKSQRLARLVLGC
ITMAYVWGKGHGDVRKVLPRNIAVPYCQLSKKLELPPILVYADCVLANWKKKDPNKPL
TYENMDVLFSFRDGDCSKGFFLVSLLVEIAAASAIKVIPTVFKAMQMQERDTLLKALLEI
ASCLEKALQVFHQIHDHVNPKAFFSVLRIYLSGWKGNPQLSDGLVYEGFWEDPKEFAG
GSAGQSSVFQCFDVLLGIQQTAGGGHAAQFLQDMRRYMPPAHRNFLCSLESNPSVREF
VLSKGDAGLREAYDACVKALVSLRSYHLQIVTKYILIPASQQPKENKTSEDPSKLEAKGT
GGTDLMNFLKTVRSTTEKSLLKEGLEHHHHHH

### Supplementary Figures

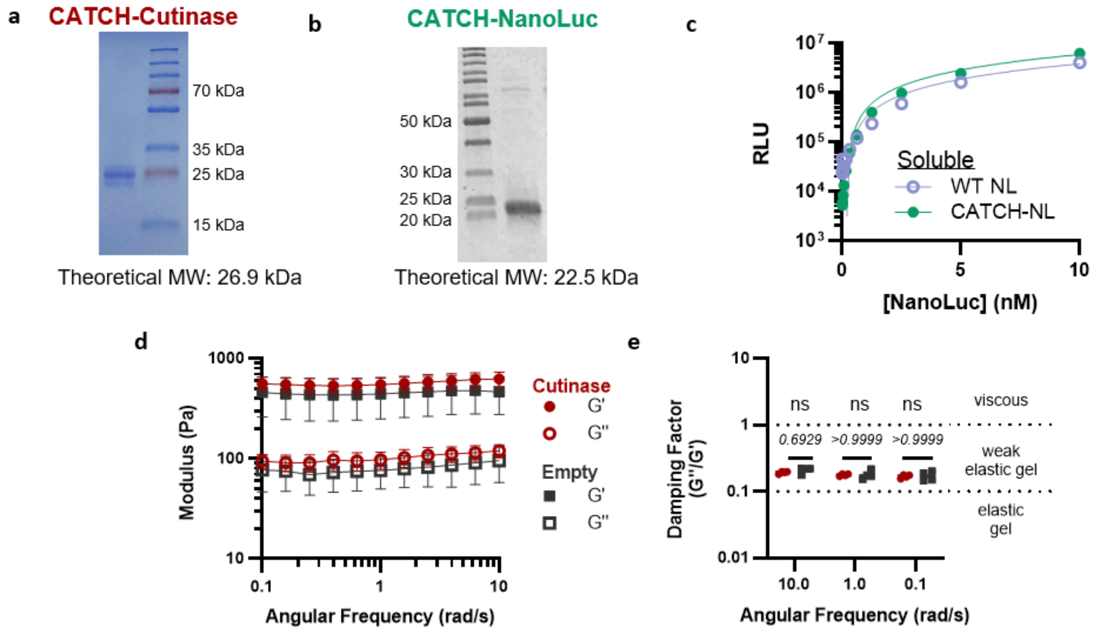


**Supplementary Figure 1: CATCH-Cutinase and CATCH-NL**

SDS-PAGE gel of (a) CATCH-cutinase and (b) CATCH-NL purified from *E coli*. (c) Luminescence output of varying concentrations wild-type (WT, light blue open circle) and CATCH-tagged NL (green circle). (d) Storage (G') and loss (G'') moduli, and (e) damping factor of CATCH-cutinase gels (red circles) and empty gels (gray squares) at 37°C after injection through a 27G needle. N = 4, ns denotes P > 0.05, ANOVA with Bonferonni post hoc. Data are presented as mean ± s.d.

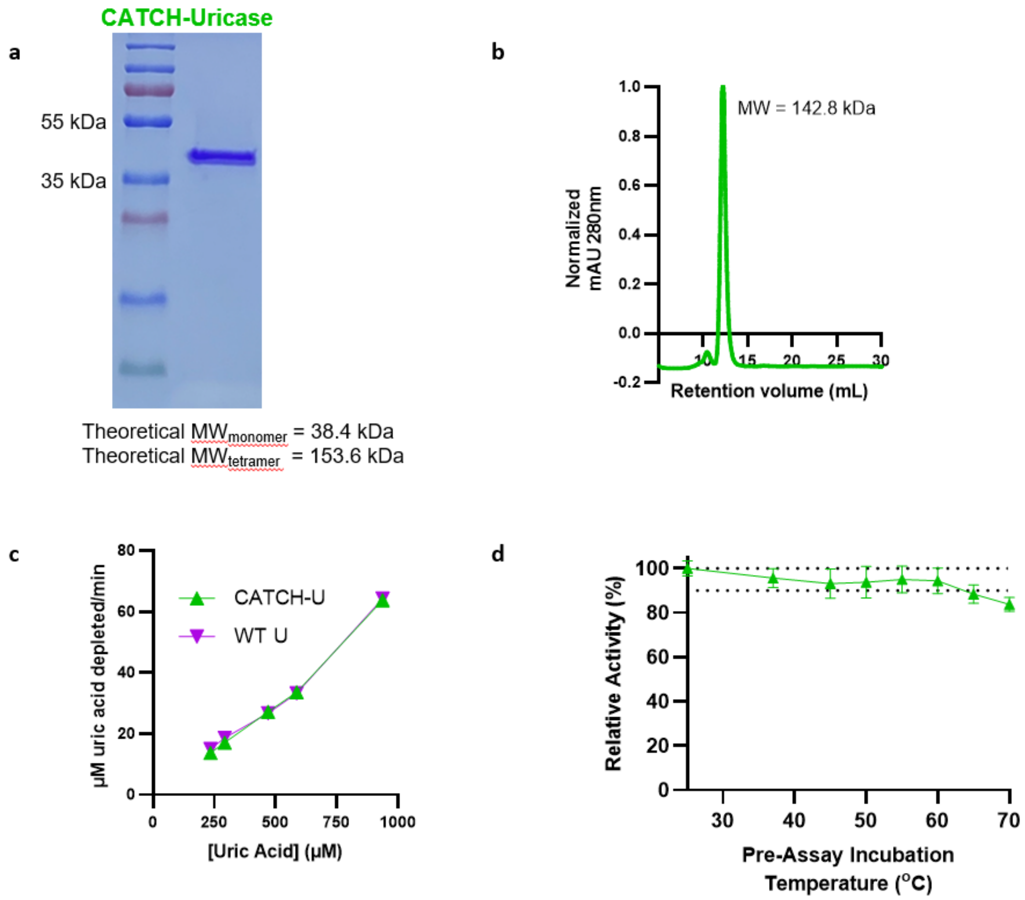


**Supplementary Figure 2: Expression and activity of CATCH-U**

(a) SDS-PAGE and (b) analytical SEC of CATCH-U purified from E. coli. (c) Velocity of uric acid depletion by WT and CATCH-U over a range of substrate concentrations (N = 2). (d) Relative activity of soluble CATCH-U after a 5-minute incubation at various temperatures, as measured by velocity of uric acid depletion normalized to velocity after room temperature incubation. Dashed lines indicate 100% and 90% activity from room temperature. (N = 2) Data are presented as mean ± s.d.

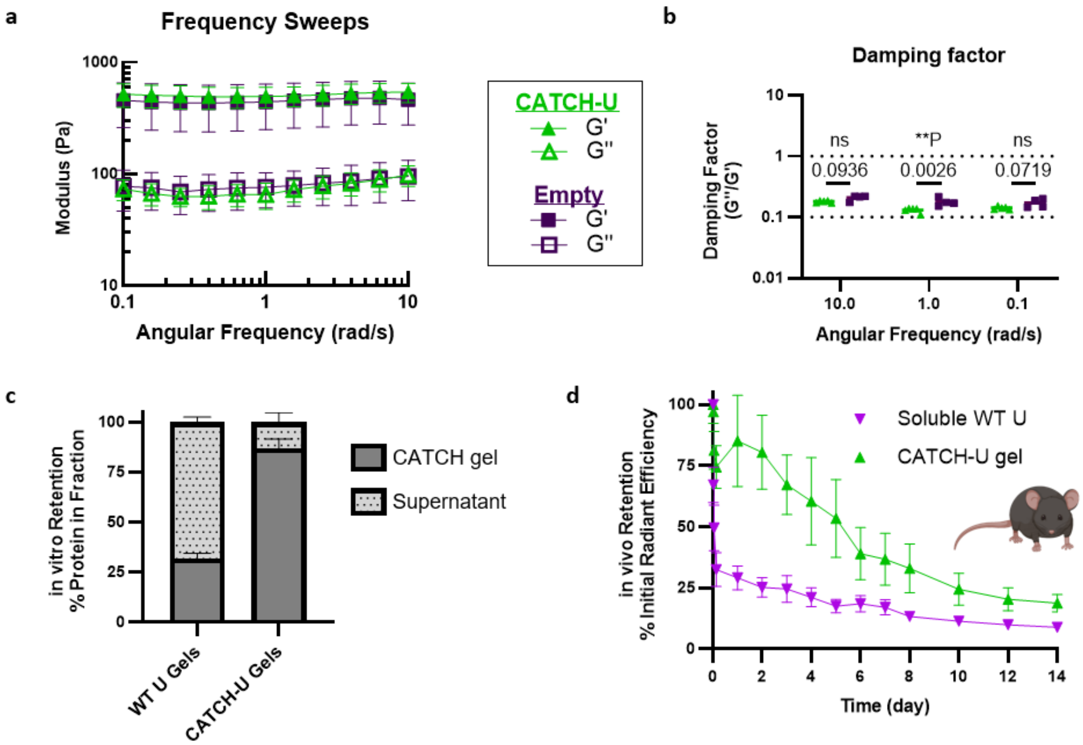


**Supplementary Figure 3: CATCH-U gel rheology and retention**

(a) Storage (G') and loss (G'') moduli, and (b) damping factor of CATCH-U (N = 5) and empty gels (N = 4) at 37$^{o}$C after injection through a 27G needle. N = 3, ns denotes $P > 0.05$, $**P < 0.01$, ANOVA with Bonferroni post hoc. (c) Percentage of WT uricase or CATCH-U retained within CATCH gels vs released into the supernatant after a 48-hour incubation at room temperature, normalized to total measured protein, N = 5. (d) Relative in vivo fluorescence of Cy5.5-labeled WT uricase (inverted purple triangle) in PBS vehicle or Cy5.5-labeled CATCH-U immobilized in CATCH gels (green triangle) subcutaneously injected into the top of the hind paw. Soluble WT uricase half-life = 0.03 d, N = 4, $R^2 = 0.915$, CATCH-U half-life = 6.9 days, N = 5, $R^2 = 0.8309$. Data are presented as mean ± s.d.

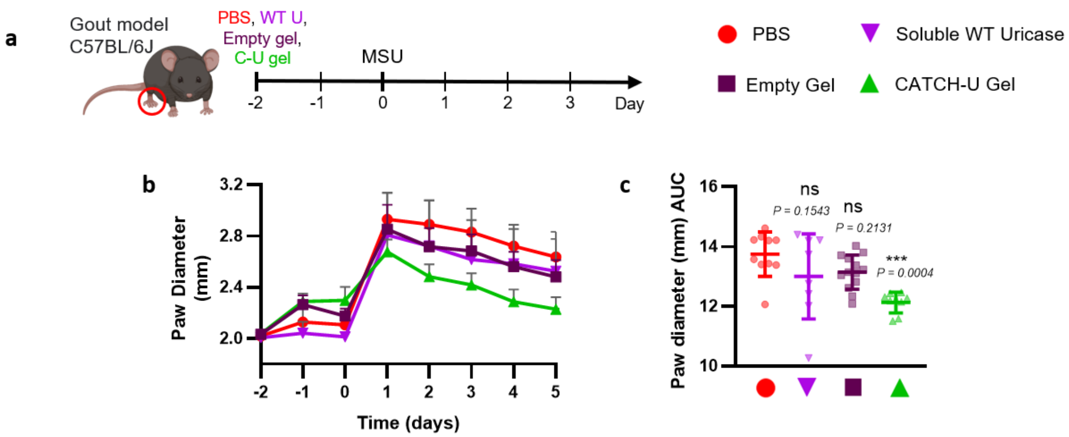


**Supplementary Figure 4: CATCH-U gel reduces paw swelling in a C57BL/6J mouse model of gouty inflammation**

(a) Schedule to evaluate inflammation following subcutaneous MSU challenge in C57BL/6J mice pretreated with PBS (red circle), soluble wild-type (WT) uricase (purple inverted triangle), empty gel (plum square), or CATCH-U gel (green triangle). (b) Raw paw caliper measurements of mouse paws and (c) area under the curve from day 0 through 5, N = 8-13, ANOVA with Dunnett's multiple comparisons test to PBS group. Data are presented as mean ± s.d.

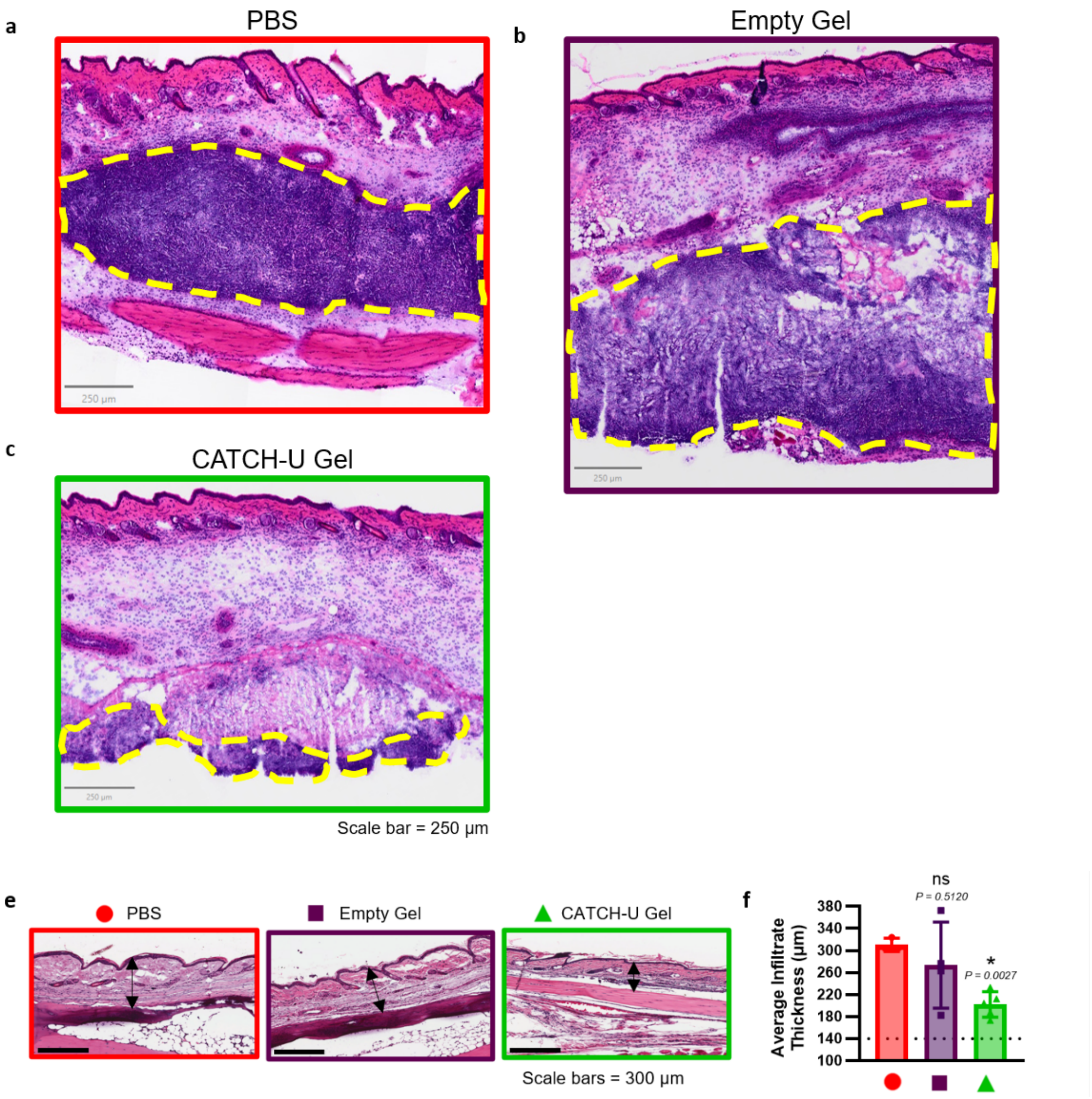


**Supplementary Figure 5: CATCH-U gel reduces immune cell infiltration in a C57BL/6J mouse model of gouty inflammation**

H&E histology on mouse paw skin 24 hours after MSU crystal challenge in mice pretreated (a) PBS, (b) empty gel, or (c) CATCH-U gel (N=1 per treatment) 48 hours prior MSU challenge. Yellow dashed line indicates MSU crystal burden and associated cell infiltration. (e) H&E histology of mouse paws 144 hours post MSU challenge. Mice received PBS (red circle), blank gels (plum square), or CATCH-U gels (green triangle) 48 hours prior to MSU crystal injection. Arrows denote cutaneous infiltrate thickness. (f) Quantification of infiltrate thickens, where the dashed line represents average paw thickness from N = 1 untreated control mouse. N = 3-5, *$P \leq 0.05$, ANOVA with Dunnett's multiple comparison test to PBS group. Data are presented as mean ± s.d.

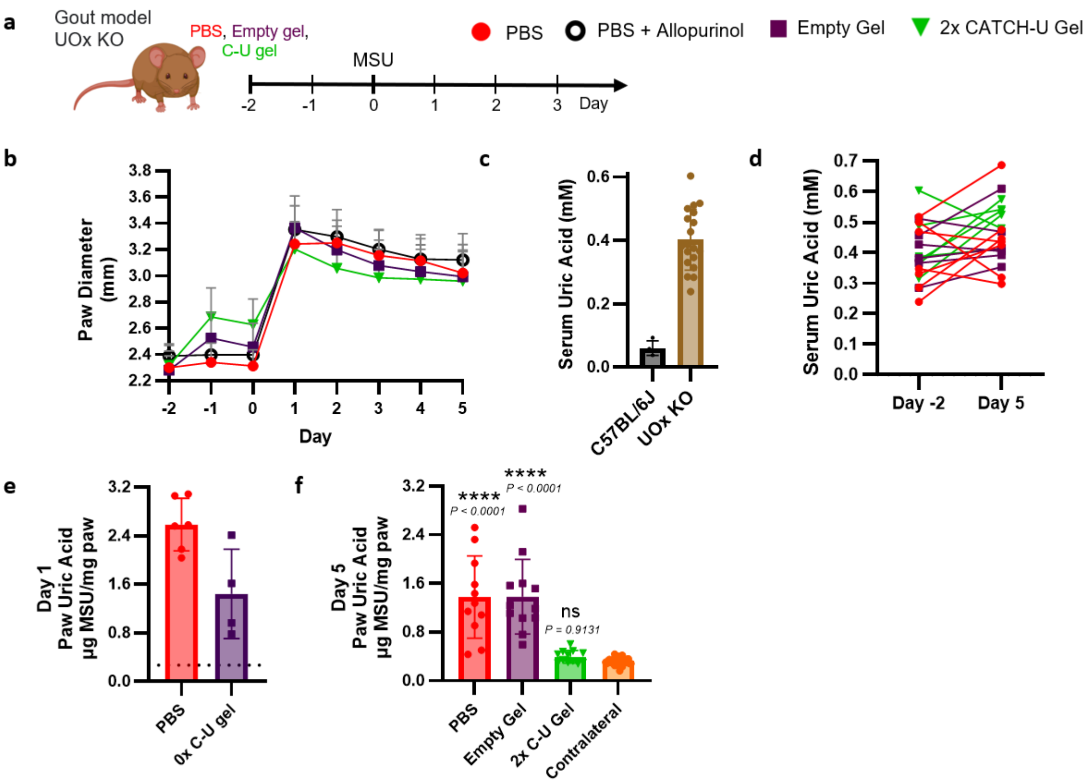


**Supplementary Figure 6: CATCH-U gels decrease gouty inflammation and MSU crystal burden in a hyperuricemic UOx KO mouse model of gout**

(a) Schedule to evaluate inflammation following subcutaneous MSU challenge in UOx KO mice pretreated with PBS (red circle), sustained on allopurinol water (black open circle), empty gel (plum square), or 2x CATCH-U gel (upside down green triangle). (b) Raw paw caliper measurements. (c) Serum uric acid of C57BL/6J mice (N=4) and UOx KO mice prior to study (N=19) and (d) at the end of the study. (e) MSU crystal burden in mouse paws 24 hours after MSU injection in mice pretreated with PBS or empty gel 48 hours prior to MSU injection, N=4-6. (f) MSU crystal burden in mouse paws 5 days after MSU injection in mice pretreated with PBS, empty gel, or 2x C-U gel, and naïve contralateral paws, N=11-12, ANOVA with Dunnett's multiple comparisons test to contralateral paws. $*P \leq 0.05$, $**P \leq 0.01$ $***P \leq 0.001$, $****P \leq 0.0001$. All data are presented as mean ± s.d.

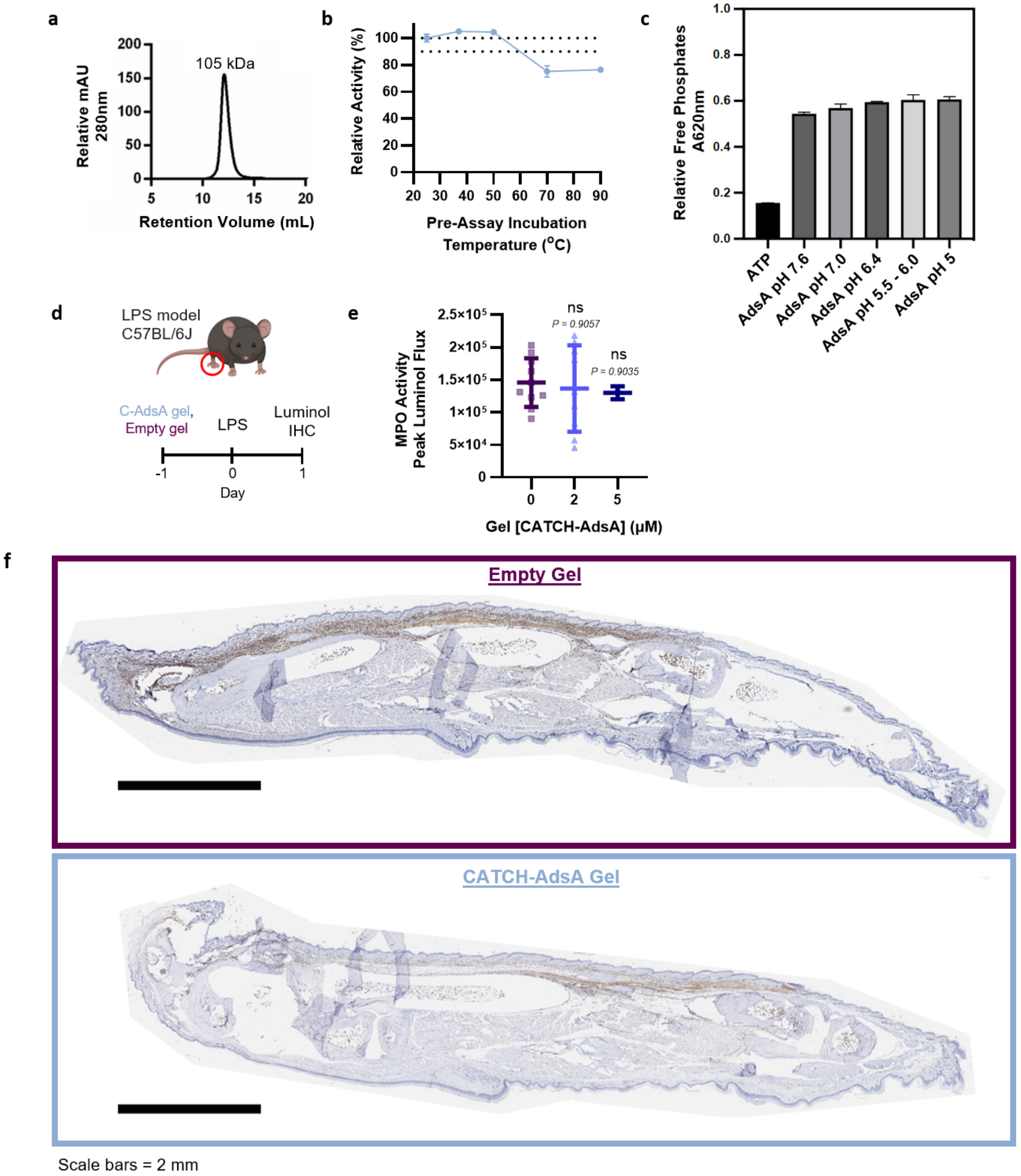


**Supplementary Figure 7: Expression and activity of soluble CATCH-AdsA; 9µM CATCH-AdsA gels decrease LPS-induced neutrophil infiltration**

(a) SEC trace of CATCH-AdsA purified from *E. coli.* (b) Relative activity of soluble CATCH-AdsA after a 5-minute incubation at the designated temperature, as measured by velocity of free phosphate generation normalized to velocity at room temperature, N=3. Dashed lines indicate 100% and 90% activity from room temperature. (c) Relative amount of free phosphates generated by soluble CATCH-AdsA over a pH range. (d) Schedule to evaluate subcutaneous LPS challenge

(Supplementary Figure 7 continued) via luminol measurements or immunohistochemistry (IHC) at t = 1 d after LPS injection. Animals received empty gels or CATCH-AdsA gels on day -1. (e) Peak MPO activity as measured by IVIS luminol bioluminescence in mice that CATCH-AdsA gels with varying concentrations of CATCH-AdsA, N = 2-13. (f) Anti-Ly6B.2 IHC of mouse hind paws at t = 1 d after subcutaneous LPS challenge in mice that received an empty gel (top, plum) or 9 μM CATCH-AdsA gel (bottom, light blue) at t = -1 d. Scale bars = 2 mm. All data are presented as mean ± s.d.

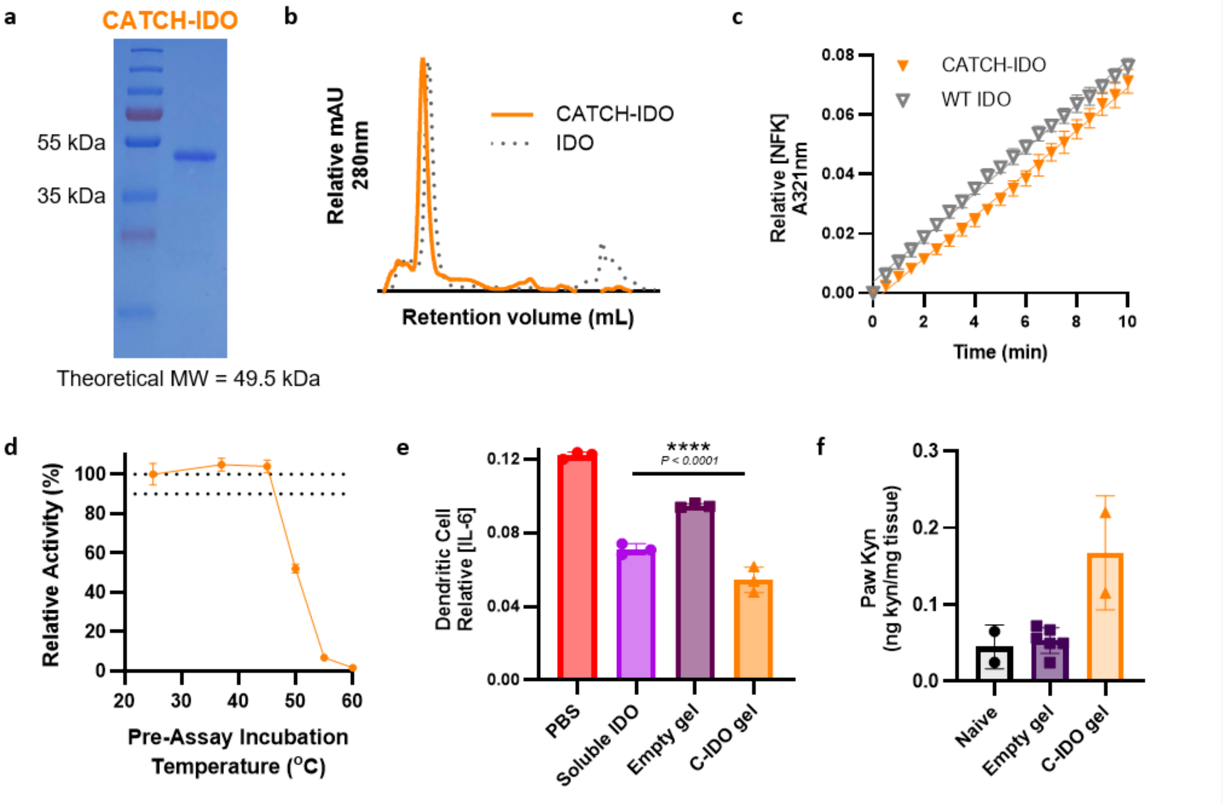


**Supplementary Figure 8: CATCH-IDO purification and activity**

(a) SDS-PAGE gel of CATCH-IDO and (b) SEC trace of CATCH-IDO and WT IDO purified from *E. coli*. (c) Relative NFK concentration produced by CATCH-IDO (orange triangle) and WT IDO (gray open triangle), N=3. (d) Relative activity of soluble CATCH-IDO after a 5-minute incubation at various temperatures, as measured by velocity of NFK production normalized to velocity after room temperature incubation. Dashed lines indicate 100% and 90% activity from room temperature, N=3. (e) Relative IL-6 produced by LPS-stimulated mouse bone marrow derived dendritic cells pretreated overnight with PBS, soluble IDO, empty gel, or CATCH-IDO gel. N=3, ****P $\leq$ 0.0001, ANOVA with Dunnett's multiple comparisons to PBS group. (f) Kynurenine levels from mouse paw tissue 1 hour after injection with empty gel or CATCH-IDO gel compared to naive tissue. N=2-6. Data are presented as mean ± s.d.

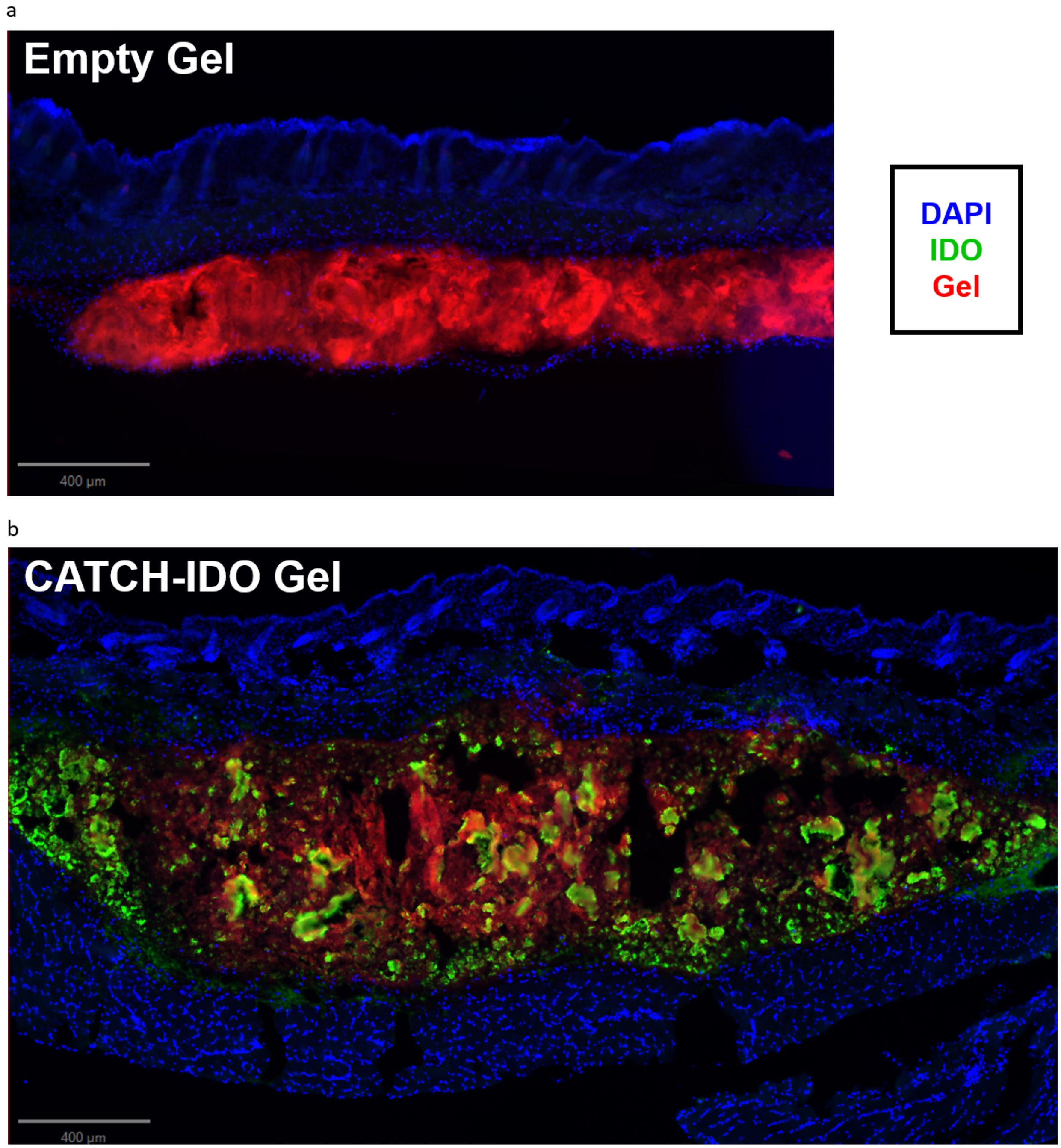


Scale bars = 400 µm

**Supplementary Figure 9: CATCH-IDO is present at a subcutaneous injection site.**

Immunofluorescent histology of mouse back skin 2 hours after injection with (a) empty gel or (b) CATCH-IDO gel (scale bar = 400µm).

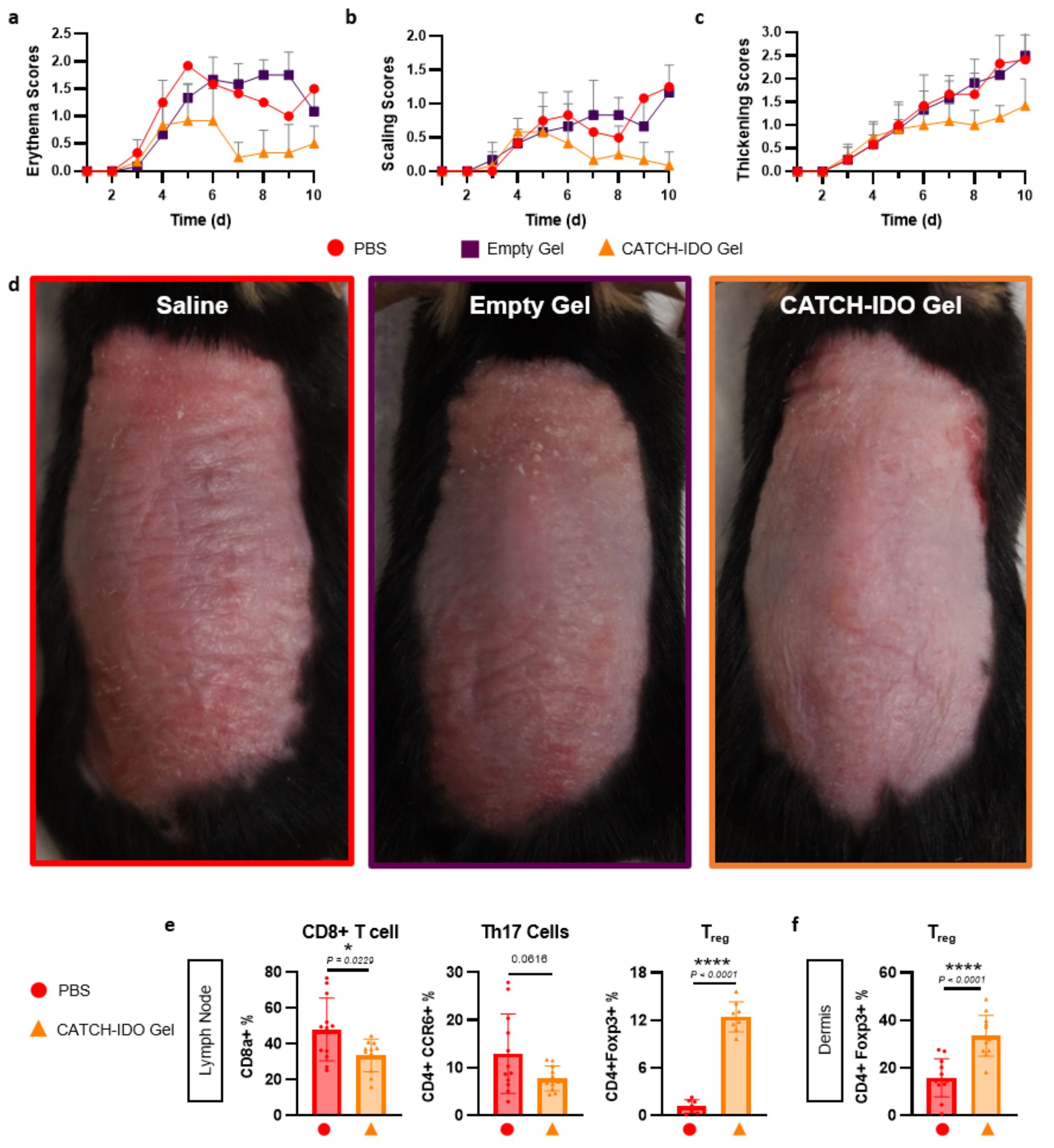


**Supplementary Figure 10: CATCH-IDO gels reduce disease severity in psoriasis**

IMQ-induced psoriasis clinical scoring for epidermal (a) erythema, (b) scaling, and (c) thickening for mice treated with PBS (red circle), empty gels (plum square) or CATCH-IDO gels (orange triangle), N=6. (d) Representative photographs of mice on day 7 of IMQ application treated with PBS (left), blank gel (middle), or CATCH-IDO gel (right) on day 3. (e) Immune cell populations in the lymph node and (f) dermis on day 7 in mice treated with PBS or CATCH-IDO gels. N=7-12, unpaired t test, *P ≤ 0.05, ****P ≤ 0.0001. All data are presented as mean ± s.d.

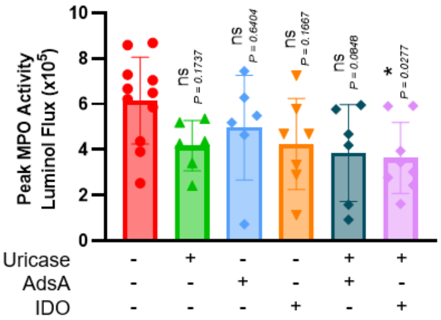


**Supplementary Figure 11: CATCH-U + CATCH-IDO dual-enzyme gels reduce neutrophil degranulation**

MPO activity as measured by luminol luminescence 24 hours after MSU challenge in C57BL/6J mice pretreated with PBS (red circle), CATCH-U gel (green triangle), AdsA gel (light blue inverted triangle), CATCH-IDO gel (orange inverted triangle), CATCH-U+AdsA gel (dark teal diamond), and CATCH-U+IDO gel (pink diamond). N=6-11, ANOVA with Dunnett's multiple comparisons to PBS, $*P \leq 0.05$. All data are presented as mean ± s.d.

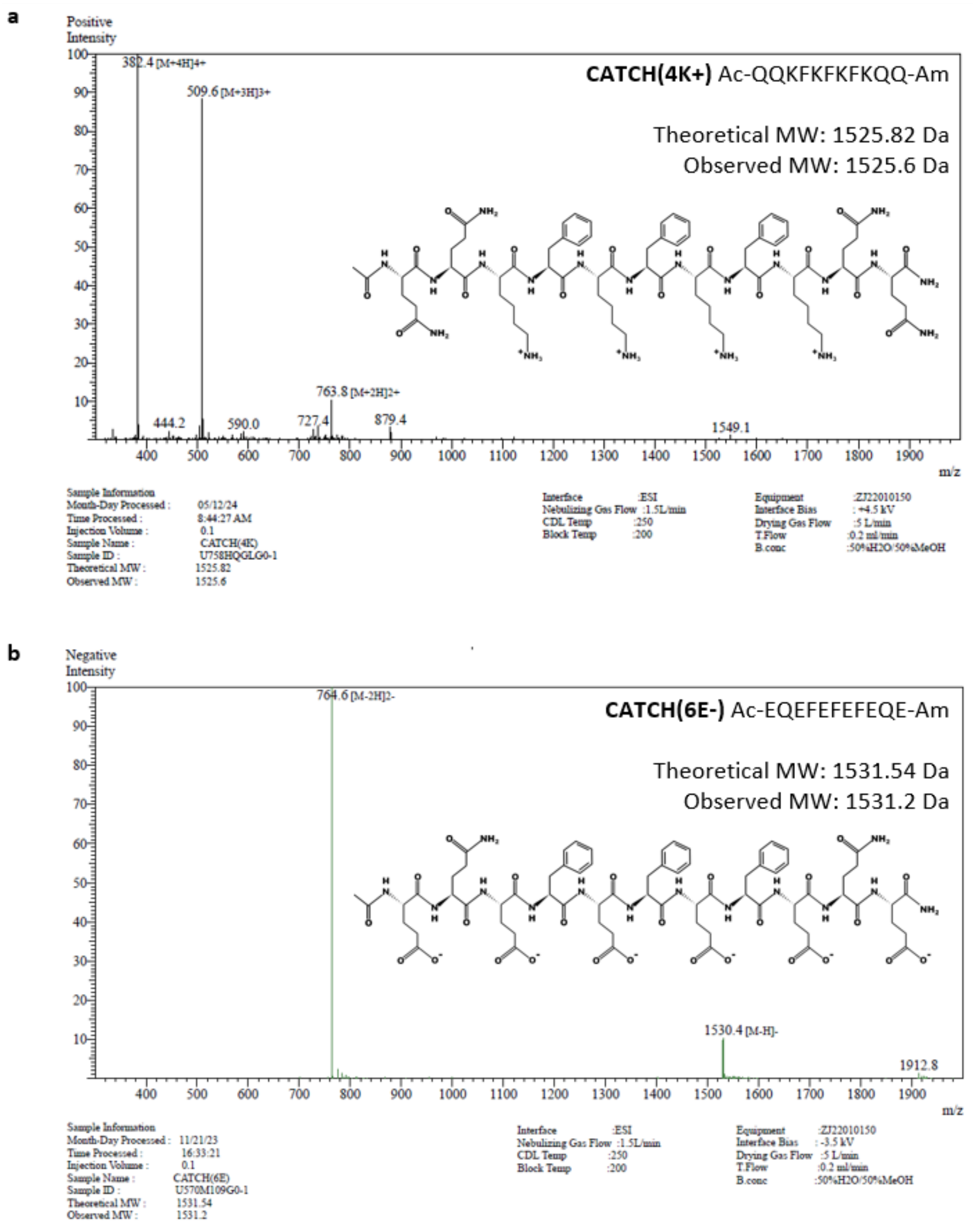


**Supplementary Figure 12: CATCH peptide chemical structures and mass spectrometry from GenScript synthesis**

The structure and observed molecular weights of (a) CATCH(+) and (b) CATCH(-)

**a**

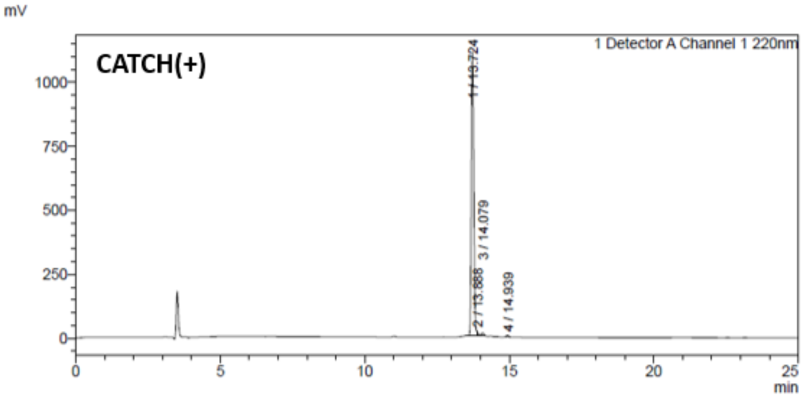


<Peak Table>

Detector A Channel 1 220nm

| Peak# | Ret. Time | Area | Height | Area% |
|---|---|---|---|---|
| 1 | 13.724 | 6571494 | 1109496 | 98.046 |
| 2 | 13.888 | 42674 | 13478 | 0.637 |
| 3 | 14.079 | 58941 | 9750 | 0.879 |
| 4 | 14.939 | 29374 | 5933 | 0.438 |
| Total | | 6702483 | 1138657 | 100.000 |

**b**

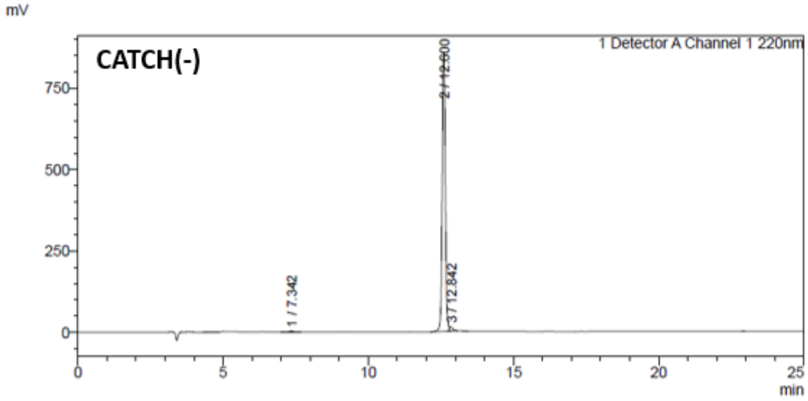


<Peak Table>

Detector A Channel 1 220nm

| Peak# | Ret. Time | Area | Height | Area% |
|---|---|---|---|---|
| 1 | 7.342 | 25206 | 3606 | 0.385 |
| 2 | 12.600 | 6434370 | 859258 | 98.261 |
| 3 | 12.842 | 88663 | 12814 | 1.354 |
| Total | | 6548239 | 875679 | 100.000 |

**Supplementary Figure 13: CATCH peptide high performance liquid chromatography chromatograms from GenScript synthesis**

(a) CATCH(+) and (b) CATCH(-) were both ≥ 98% pure
